# Cadherin 16 promotes sensory gating via the endocrine corpuscles of Stannius

**DOI:** 10.1101/2024.09.23.614609

**Authors:** Susannah S. Schloss, Zackary Q. Marshall, Nicholas J. Santistevan, Stefani Gjorcheska, Amanda Stenzel, Lindsey Barske, Jessica C. Nelson

**Affiliations:** Department of Cell and Developmental Biology; University of Colorado Anschutz Medical Campus School of Medicine, Aurora, CO, USA; Division of Human Genetics, Cincinnati Children’s Hospital Medical Center, Department of Pediatrics, University of Cincinnati College of Medicine, Cincinnati, OH, USA

**Keywords:** zebrafish, sensorimotor gating, calcium homeostasis, *cdh16*, *stc1l*, *pappaa*, corpuscles of Stannius, habituation, acoustic startle

## Abstract

Sensory thresholds enable animals to regulate their behavioral responses to environmental threats. Despite the importance of sensory thresholds for animal behavior and human health, we do not yet have a full appreciation of the underlying molecular-genetic and circuit mechanisms. The larval zebrafish acoustic startle response provides a powerful system to identify molecular mechanisms underlying establishment of sensory thresholds and plasticity of thresholds through mechanisms like habituation. Using this system, we identify Cadherin 16 as a previously undescribed regulator of sensory gating. We demonstrate that Cadherin 16 regulates sensory thresholds via an endocrine organ, the corpuscle of Stannius (CS), which is essential in zebrafish for regulating Ca^2+^ homeostasis. We further show that Cadherin 16 regulates whole-body calcium and ultimately behavior through the hormone Stanniocalcin 1L, and the IGF-regulatory metalloprotease, Papp-aa. Finally, we demonstrate the importance of the CS through ablation experiments that reveal its role in promoting normal acoustic sensory gating. Together, our results uncover a previously undescribed brain non-autonomous pathway for the regulation of behavior and establish Ca^2+^ homeostasis as a critical process underlying sensory gating *in vivo*.

## Introduction

Animals use sensory cues to evade threats in the environment. The acoustic startle response provides a crucial defensive mechanism, observed in species throughout the animal kingdom^1,2^. Although it is critical that animals be able to mount escape responses to threatening stimuli, they must also be able to distinguish between threatening and non-threatening stimuli. Sensory thresholds enable animals to make this distinction, dictating the minimum stimulus intensity that elicits a response^3–5^. In the case of the acoustic startle response in larval zebrafish, sensory thresholds are established during development and differ between animals, representing a form of behavioral individuality^6^. Moreover, thresholds established during development can be transiently modified through plasticity mechanisms like habituation^7,8^. In humans, a variety of neurological disorders, including schizophrenia, autism spectrum disorder, and migraine, are associated with differences in the ability to properly threshold or habituate to sensory stimuli^9,10^. Therefore, understanding the underlying biology may shed light on molecular mechanisms underlying disease.

Previous work has identified multiple molecular pathways that regulate sensory gating in larval zebrafish^3,4,11–15^. Many of these molecular regulators are expressed in or affect the activity of cells comprising the acoustic startle circuit, including the IGF-regulatory metalloprotease *pappaa*^11^, the voltage-gated K^+^ channel subunit *kcna1a*^14^, the palmitoyltransferase *hip14*^14^, the cytoskeletal regulator *cyfip2*^3^, and the Ca^2+^-sensing receptor *casr*^12,16^. Prior work has also probed key neurotransmitter signaling pathways that regulate acoustic startle response gating^7,8,17^. Together, while this work has placed molecular mechanisms of behavior in the context of circuit function, most of the identified mechanisms function autonomously in the brain. How brain non-autonomous regulators of internal state, including whole-body homeostatic states might contribute is thus far largely unexplored.

In this study, we identify a novel brain non-autonomous mechanism key for promoting sensory gating. We find that Cadherin 16 (encoded by *cdh16*) functions in the pronephros-derived corpuscles of Stannius to regulate Ca^2+^ homeostasis and ultimately sensory thresholds in larval zebrafish. This system provides an ideal model for understanding how Ca^2+^ homeostasis regulates sensory thresholds. In zebrafish, the corpuscles of Stannius produce the Ca^2+^-regulatory hormone Stanniocalcin 1L^18,19^. Stanniocalcin 1L then functions to limit the proliferation and function of epithelial cells called ionocytes, which are specialized for Ca^2+^ uptake^20^. In particular, a specific class of ionocytes, termed Na+/H+-ATPase-rich (NaR) cells, promote Ca^2+^ uptake from the environment^21^. Stanniocalcin 1L limits their proliferation through suppression of a metalloprotease, Papp-aa, expressed in NaR ionocytes^20^. Consequently, zebrafish *pappaa* loss-of-function mutants show reduced bone calcification^22^. PAPP-A is similarly crucial for calcium homeostasis in mammals. Homozygous loss-of-function mutations in *PAPPA2* in humans are associated with growth deficits and reduced bone mineralization^23^. Interestingly, Papp-aa has also been identified as a key regulator of acoustic and visual behaviors in zebrafish^11^.

Here we find that this Ca^2+^-regulatory pathway functions in the context of sensory gating. Through genetic epistasis, we find that Cdh16 functions through Stanniocalcin 1L and ultimately Papp-aa to regulate whole-body Ca^2+^, which in turn broadly regulates behavioral thresholds, with opposite impacts on visually and acoustically evoked startle responses. Therefore, our results highlight a link between Papp-aa and Cdh16 function and underscore a crucial role for Ca^2+^ homeostasis in the regulation of sensory gating and behavior. Interestingly, human patient data also support a crucial role for Ca^2+^ homeostasis in the regulation of sensory gating: hypocalcemia in human patients is associated with seizures and psychotic symptoms, including auditory hallucinations^24,25^.

## Results

### Cadherin 16 regulates acoustic startle response thresholds and habituation learning

At 5 days post-fertilization (dpf), larval zebrafish respond to threatening acoustic stimuli via a short-latency acoustic startle response, or short-latency C-bend (SLC)^2,26^. At this stage, zebrafish are also capable of distinguishing between high-intensity stimuli that necessitate a fast response, and lower intensity stimuli that often receive a response in the form of a long-latency C-bend or are ignored entirely^3,11,26^. Sensorimotor gating mechanisms, including the developmental establishment of acoustic startle response thresholds, enable animals to make these distinctions between threatening and non-threatening acoustic stimuli^3^. Moreover, thresholds established during development can be transiently modified in 5 dpf larvae through plasticity mechanisms like habituation^7,8,11^.

Through a forward genetic screen, a large collection of molecular regulators of sensorimotor gating were identified, including genes regulating (1) initial establishment of acoustic startle response thresholds, (2) plasticity of thresholds through habituation, and (3) the decision to perform a short-latency versus a long-latency C-bend^3,11–15^. *irresistible^p173^* mutants were identified based on their hypersensitivity to acoustic stimuli and inability to modulate response frequency through habituation^11^. To quantify these phenotypes, we exposed *irresistible* mutants to a series of acoustic stimuli, ranging in intensity from 0.54g to 51.1g as previously described (see methods)^3,4^. *irresistible* mutants exhibit an increased sensitivity to acoustic stimuli, responding at higher rates than their siblings across multiple stimulus intensities, indicating deficits in sensory gating **(Fig 1A)**. Next, we measured habituation by presenting animals with 40 high-intensity (51.1g) stimuli, each separated by a 3-second inter-stimulus interval (ISI). We found that *irresistible* mutants continue to respond at a high rate throughout the habituation assay, indicating deficits in the ability to dynamically tune acoustic startle response thresholds (**Fig 1B-C**).

**Figure 1.**
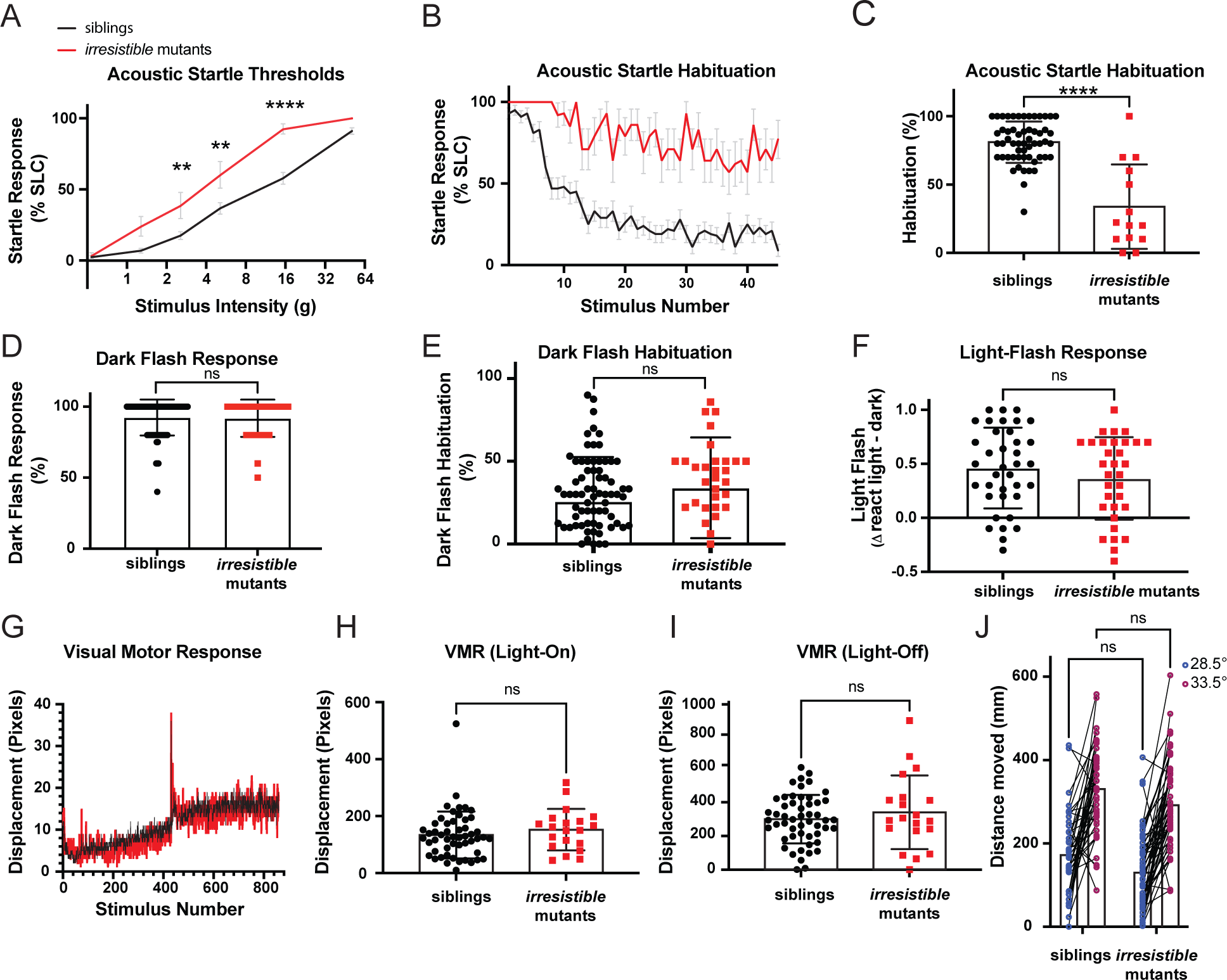
*irresistible* mutations suppress habituation and cause hypersensitivity to acoustic stimuli. **A)** *irresistible* mutants (n=13) display heightened sensitivity to acoustic stimuli as compared to heterozygous and wild type (WT) siblings (n=52). Error bars show SEM. Differences in startle sensitivity were calculated using a two-way ANOVA with a Šídák’s multiple comparisons test (**p<0.01, ****p<0.0001). **B)** *irresistible* mutants (n=14) fail to habituate to repeated acoustic stimuli when compared to siblings (n=58), error bars show SEM. **C)** *irresistible* mutants (n=14) have lower habituation (****p<0.0001, Mann-Whitney test) in relation to their siblings (n=56). Error bars show SD. **D)** *irresistible* mutants (n=33) and siblings (n=80) have no difference (p=0.8615, Mann-Whitney test) in their response to dark flash stimuli. Error bars show SD. **E)** *irresistible* mutants (n=33) and siblings (n=80) display no differences in habituation to dark flash stimuli (p=0.0686, Mann-Whitney test). Error bars show SD. **F)** *irresistible* mutants (n=32) have no differences (p=0.2983, unpaired t-test) in light flash reactivity as compared to their siblings (n=37). Error bars show SD. **G)** *irresistible* mutants (n=20) display normal visual motor (VMR) behaviors relative to their siblings (n=52). **H)** *irresistible* mutants (n=20) and siblings (n=52) display no difference (p=0.2471, Mann-Whitney test) in their responses to whole field illumination in VMR assay. **I)** *irresistible* mutants (n=20) and siblings (n=52) do not show significantly different responses to whole field loss-of-illumination in VMR assay (p=0.7223, Mann-Whitney test). Error bars show SD. J) *irresistible* mutants (n=54) and siblings (n=42) have no significant differences in their movement at baseline temperature (p=0.0877, two-way ANOVA with Šídák’s multiple comparisons test) and both respond to high temperature with increased locomotion (difference between mutants and siblings: p=0.1231, two-way ANOVA with Šídák’s multiple comparisons test).

Despite the dramatic impacts on their ability to threshold acoustic stimuli, *irresistible* mutants are adult-viable and fertile. To examine whether *irresistible* specifically regulates acoustic sensory gating, or has broader effects, we tested visual startle response rates (O-bend responses to whole-field loss of illumination or dark flash)^27^ **(Fig 1D)**, habituation to dark flash stimuli^7,28,29^ **(Fig 1E)**, light flash responses^27^ **(Fig 1F)**, visuomotor responses^30^ **(Fig 1G-I)**, and ability to respond to thermal stimuli^31^ **(Fig 1J)**. We found no significant differences between *irresistible* mutants and their siblings, consistent with a specific deficit in developmental and acute regulation of acoustic thresholds in mutant larvae.

To map the genetic locus responsible for the *irresistible* phenotype, we conducted whole-genome sequencing followed by homozygosity mapping as previously described^11^. This uncovered a premature stop codon (Y657*) in the *cdh16* gene, encoding the calcium-dependent cell-adhesion protein, Cadherin 16. Like other members of the 7D family of cadherins, Cadherin 16 has 7 extracellular cadherin domains, a transmembrane domain (TM), and a short intracellular domain^32,33^. Y657* results in a termination codon after the 6^th^ Cadherin domain, prior to the TM domain **(Fig 2A)**. To determine whether the premature stop codon in the *cdh16* locus is causal for the acoustic hypersensitivity and habituation phenotypes, we used CRISPR-Cas9 genome editing to generate an independent loss-of-function allele, *co79*, in *cdh16* in otherwise wild type animals. *co79* results in a 10bp deletion in exon 2, resulting in a frameshift and premature stop codon (F38del[SPSCQISL*]FSX8) **(Fig 2A)**. Like *p173*, animals homozygous for the *co79* mutant allele are hypersensitive to acoustic stimuli and fail to habituate (**Fig 2B-D**). Conversely, habituation and startle sensitivity assays demonstrated that animals heterozygous for either *co79* or *p173* are indistinguishable from their wild type siblings on these measures **(Fig 2E-G)**. Finally, through complementation testing, we determined that larvae carrying a combination of both mutant alleles (*cdh16^p173/co79^*) fail to habituate and exhibit hypersensitivity to low-intensity acoustic stimuli **(Fig 2E-G)**. Together, these data demonstrate that Cadherin 16 regulates the establishment and dynamic tuning of acoustic startle thresholds through habituation.

**Figure 2.**
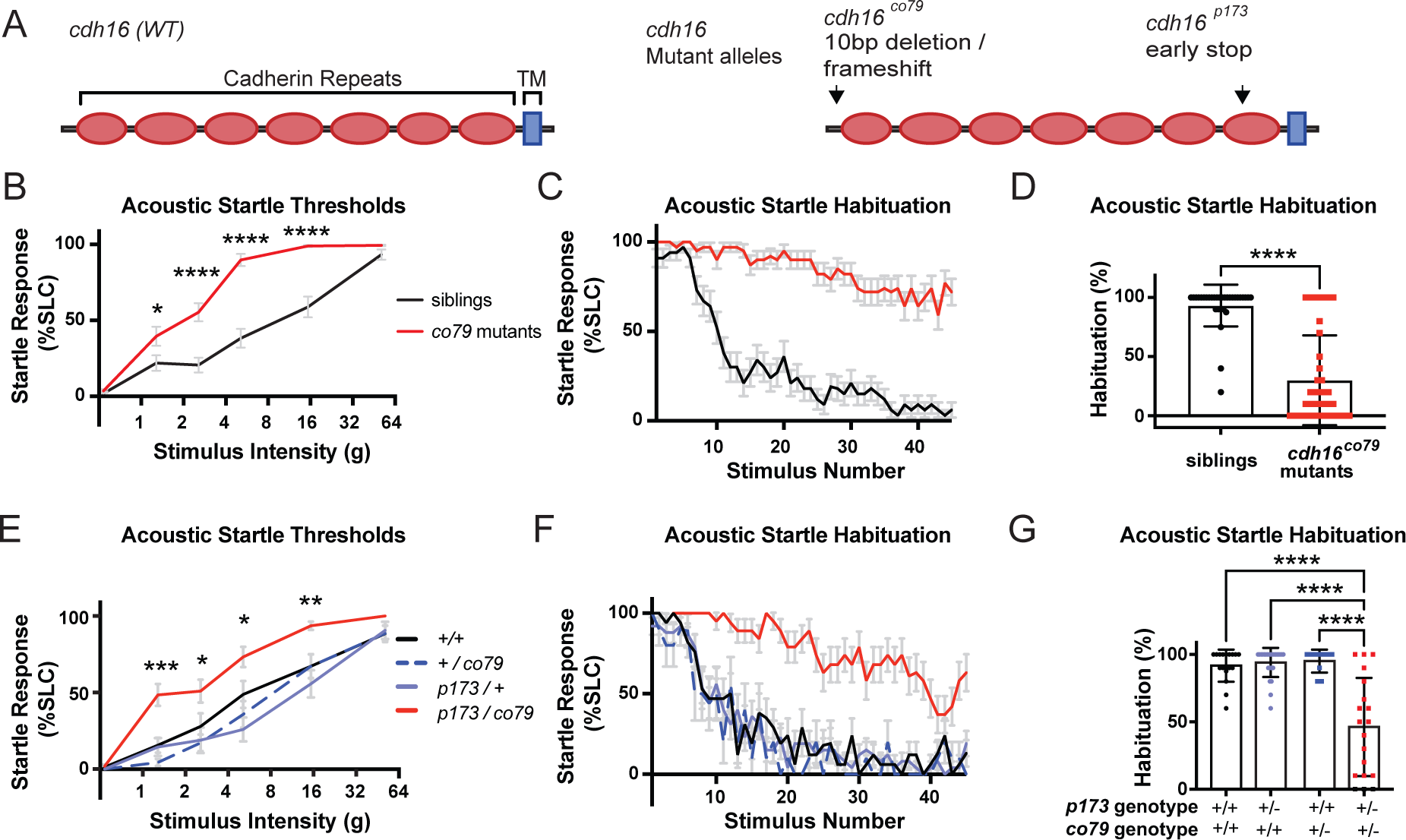
*irresistible^p173^* is an allele of the Cadherin-encoding gene *cdh16*. **A)** Conceptual translation of *cdh16,* the predicted consequences of the *irresistible^p173^*, and *cdh16^co79^* mutations. **B)** *cdh16^co79^* mutants (n=39) have decreased thresholds to low intensity acoustic stimuli as compared to their siblings (n=33) (*p=0.0319, ****p<0.0001, two-way ANOVA with Šídák’s multiple comparisons test). **C)** *cdh16^co79^* mutants (n=39) continue responding to repeated acoustic stimuli while their siblings (n=33) habituate. **D)** *cdh16^co79^* mutants (n=39) have significantly impaired habituation (****p<0.0001, Mann-Whitney test) compared to siblings (n=32). **E)** *cdh16^p173^* / *cdh16^co79^* transheterozygotes (n=22) have increased sensitivity to acoustic stimuli when compared to *cdh16^p173^* heterozygotes (n=17), *cdh16^co79^* heterozygotes (n=14), and wild types (n=16). A two-way ANOVA with Tukey’s multiple comparisons test was used to calculate the difference in SLC% between all groups. Differences between *cdh16^p173/co79^* vs. WT (+/+) are represented with p values on the plot: ***p=0.0005, for the difference between WT and transheterozygotes, *p<0.03, **p=0.0086. **F)** *cdh16^p173^* / *cdh16^co79^* transheterozygotes (n=19) fail to habituate to high intensity acoustic stimuli while wild type (n=17), *cdh16^co79^* heterozygotes (n=10), and *cdh16^p173^* heterozygotes (n=26) habituate normally. **G)** *cdh16^p173^* / *cdh16^co79^* transheterozygotes (n=19) have significantly lower habituation percentages p<0.0001 compared to wild types (n=17), *cdh16^co79^* heterozygotes (n=10), and *cdh16^p173^* heterozygotes (n=25). Differences in habituation between groups were calculated using a two-way ANOVA with Tukey’s multiple comparisons test. Error bars in **B, C, E**, and **F** indicate SEM. Error bars in **D** and **G** indicate SD.

### Cadherin 16 expression is sufficient after the development of the acoustic startle circuit to restore acoustic startle thresholds and habituation

The neuronal circuits required for the performance of the acoustic startle response are in place by 4dpf. By this stage, animals reliably perform acoustic startle responses to high-intensity stimuli and exhibit robust habituation learning^2,34^. Cadherin proteins regulate many developmental processes throughout the body, including the assembly of neuronal circuits^35^. Therefore, we wondered whether *cdh16* is required for the assembly of the acoustic startle circuit, or whether it might be required for the maintenance, function, or maturation of the acoustic startle circuit. To test this, we generated a transgene expressing *cdh16* under the control of the *hsp70* heat-shock activated promoter. We found that ubiquitous, heat-shock induced expression of *cdh16* at 3 and 4dpf rescued acoustic startle thresholds and habituation at 5dpf, consistent with a role for Cdh16 during development. The same manipulation had no significant effect in sibling animals overexpressing *cdh16* **(Fig 3A-B)**. However, expression of *cdh16* at 2 and 3dpf did not restore normal behavior measured at 5dpf, suggesting that maintenance of Cadherin 16 expression at the time of behavior testing is required for the regulation of sensory-evoked behaviors **(Fig 3C-D)**. Moreover, we determined that induced expression of *cdh16* at 4 and 5dpf rescued acoustic startle thresholds and habituation measured at 6dpf, consistent with a role for Cdh16 in the regulation of sensory processing after the establishment of the acoustic startle circuit **(Fig 3E-F)**. To broadly examine how *cdh16* might impact neuronal development, we performed whole-brain morphometric analyses in *cdh16* mutants versus siblings across 294 molecularly-defined brain regions^36,37^. This unbiased approach for assessing brain development revealed minimal changes in size across these regions **(Fig 3G, Supplemental Fig 1)** consistent with our rescue data and underscoring a role for Cdh16 in regulating nervous system function rather than early nervous system development.

**Figure 3.**
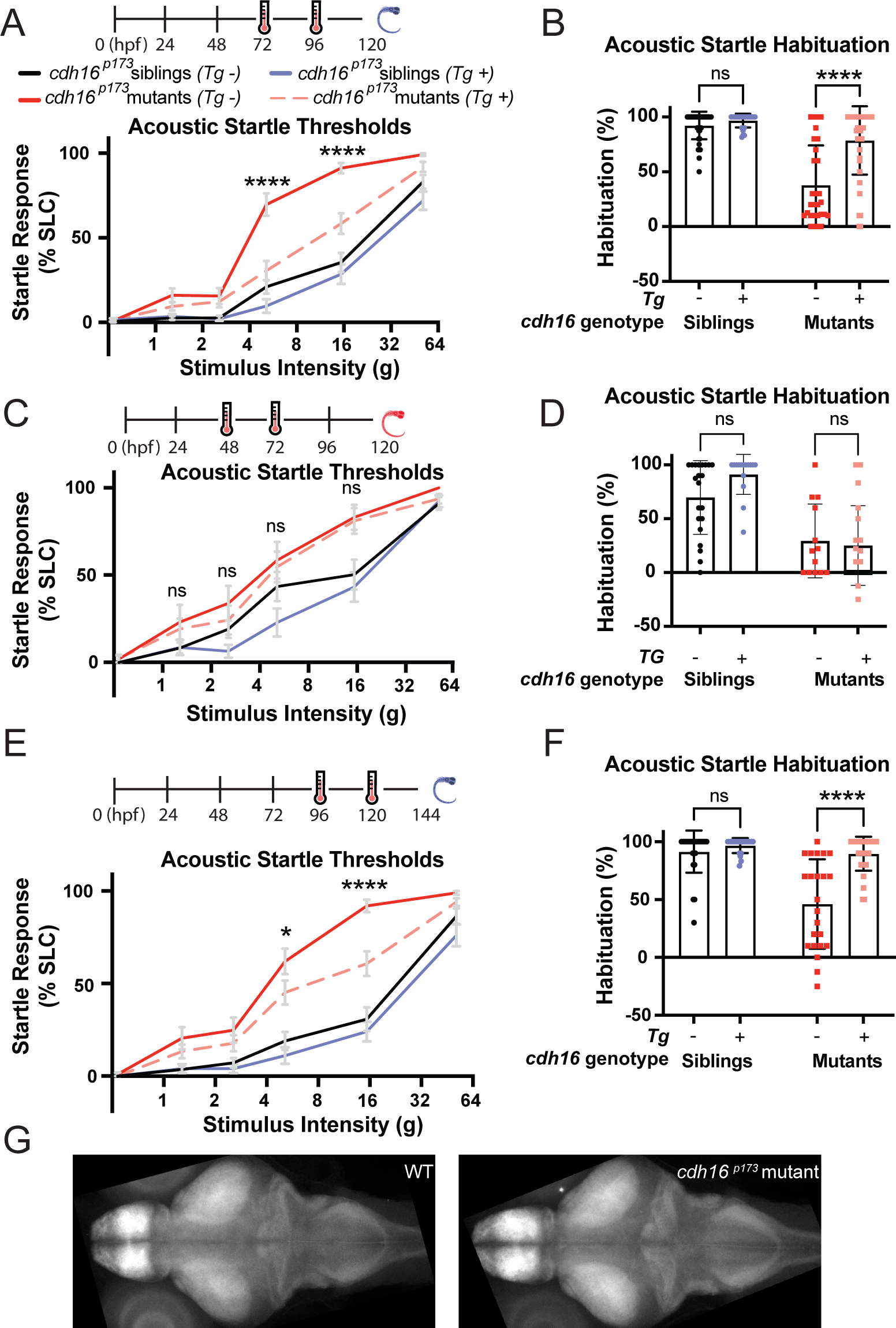
Ubiquitous expression of *cdh16* after circuit development restores habituation and acoustic sensitivity. **A)** *hsp70p:cdh16-p2a-mKate* expression was induced at 72 and 96hpf (hours post-fertilization) via heat-shock. Behavior testing and analysis performed at 120hpf. Induction of *cdh16* expression in *cdh16^p173^* mutants (n=38) results in significantly lower startle sensitivity compared to *cdh16^p173^* mutants that are heat-shocked but do not carry the transgene (n=25). ****p<0.0001. **B)** Heat-shock as in **A** has no effect on habituation (p=0.7204) of siblings (n=41 with the transgene versus n=26 without). In contrast, heat-shock induction of *cdh16* expression significantly restores habituation (p<0.0001) in *cdh16^p173^* mutants carrying the transgene (n=38) in comparison to transgene negative mutants (n=29). **C)** *hsp70p:cdh16-p2a-mKate* expression was induced at 48 and 72hpf via heat-shock. Behavior testing and analysis performed at 120hpf. Acoustic startle sensitivity is not significantly restored in *cdh16^p173^* mutants carrying the heat-shock transgene (n=19) when compared to mutants with no transgene (n=13). These data are consistent with a requirement for maintenance of *cdh16* expression during behavior, (p>0.7 for all stimulus intensities.) **D)** Heat-shock as in **C** has no effect on habituation (p=0.1073) in siblings expressing the transgene (n=15) in relation to siblings not expressing the transgene (n=21). Similarly, the difference in acoustic startle habituation in transgene-expressing mutants (n=19) and mutants not expressing the transgene (n=13) is not significant (p=0.9199). **E)** *hsp70p:cdh16-p2a-mKate* expression was induced at 96 and 120hpf (after the acoustic startle circuit is functional) via heat-shock. Behavior testing and analysis performed at 144hpf. Hypersensitivity is rescued in *cdh16^p173^* mutants (n = 27) carrying the transgene as compared to mutants lacking the transgene (n=20). (*p=0.0380, **** p<0.0001). **F)** Heat-shock as in E has no effect on acoustic startle habituation (p=0.6607) in siblings carrying the transgene (n=21) compared to siblings lacking the transgene (n=28). Conversely, *cdh16* expression restores habituation to acoustic stimuli (p<0.0001) in mutants carrying the transgene (n=31) as compared to mutants without the transgene (n=23). For **A, C, and E,** error bars indicate SEM. For **B, D, and F**, error bars indicate SD. **G)** Representative whole-brain stacks for WT (left) and cdh16^p173^ mutants (right), showing a lack of brain volume changes at 6dpf.

### *cdh16* is expressed in the corpuscles of Stannius

In mammals, *cdh16* is primarily expressed in the kidney and the thyroid^38,39^. In zebrafish, while *cdh16* expression in the brain has been documented at 10 days post-fertilization^40^, others have found that at earlier stages *cdh16* is primarily expressed in the developing pronephros or embryonic kidney^41^. Around 2 days post-fertilization, *cdh16* expression becomes largely restricted to an endocrine organ called the corpuscle of Stannius (CS), which is extruded from the pronephros and secretes Stanniocalcin 1, a calcium-regulatory hormone^41^. Given our finding that *cdh16* is required for sensory gating after 2dpf, we wondered whether *cdh16* expression might persist in the CS beyond this early developmental timepoint. To address this question, we used *in situ* hybridization chain reaction (HCR), examining *cdh16* expression in whole-mount embryos and larvae from 24 hours post-fertilization (hpf) through 144hpf (**Fig 4A-G**). At all time points, we found that *cdh16* was strongly expressed either in the pronephros (24hpf, **Fig 4B**) or the CS (48-144hpf) **(Fig 4C-G)**. From these data, we predicted that CDH16 might be required outside the brain to regulate acoustic startle thresholds.

**Figure 4.**
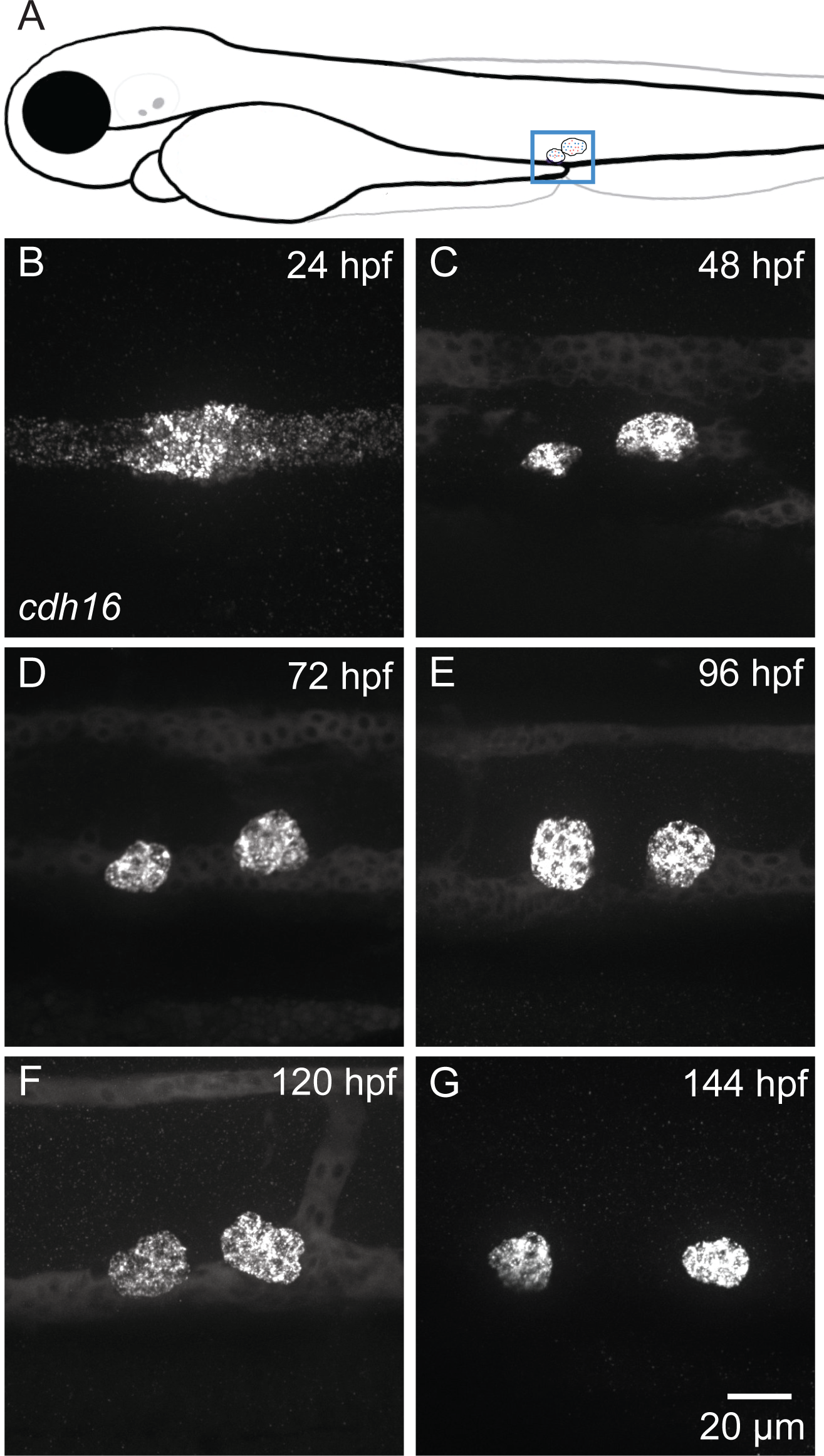
*cdh16* is expressed in the corpuscles of Stannius (CS) during embryonic and larval development. **A)** Schematic indicating the position of the corpuscles of Stannius (blue box) in the context of the whole larva. (**B-G)** Whole-mount *in situ* hybridization chain reaction (HCR) using probes against *cdh16.* Maximum projections of confocal stacks show the larval zebrafish pronephros **(B)** and corpuscles **(C-G)**. **B)** *cdh16* puncta are enriched in distal pronephros where the CS will be extruded. **(C-F)** *cdh16* signal is present in the CS and kidney from 48hpf to 120hpf. **G)** By 144hpf *cdh16* signal is present in the CS but is no longer detectable in the kidney. Shown are representative images, n=5 larvae were imaged per timepoint.

### Cadherin 16 promotes the function of PAPP-AA through the regulation of the hormone Stanniocalcin 1L

Morpholino knockdown of *cdh16* in embryonic zebrafish leads to a dramatic increase in the expression of the *stanniocalcin 1l* gene encoding the hormone, Stanniocalcin^41^. In zebrafish and mammals, Stanniocalcin 1 inhibits the metalloprotease PAPP-AA^20,22,42,43^, which is a known regulator of acoustic startle sensitivity and habituation in larval zebrafish^11^. *pappaa* mutants largely phenocopy *cdh16* with one exception: *pappaa* mutants are not responsive to dark-flash, or whole-field loss of illumination^11,44^. We hypothesized that excessive *stc1l* expression in *cdh16* mutants inhibits *pappaa,* precluding appropriate acoustic startle thresholding and plasticity of thresholds through habituation. To test our hypothesis, we set out to confirm that *cdh16* mutants, like *cdh16* morphants, show increased expression of *stc1l*. We found that as in *cdh16* morphants, *stc1l* expression was strongly increased in *cdh16* loss-of-function mutants (**Fig 5A**). Next, we wondered whether loss of *cdh16* might lead to a change in *stc1l* expression in the brain. To test this, we dissected 5dpf larval zebrafish, separating the trunk and the head, and performed RT-qPCR in each tissue independently in mutants and siblings. We found that while *stc1l* was strongly upregulated in the trunk (which contains the CS), (**Fig. 5B**) there was no change in the head (**Fig. 5C**), consistent with a CS-specific role of Cdh16 in regulating *stc1l* expression. Finally, to probe for a role for Cdh16 in neurons, we used the Gal4/UAS system to express *cdh16* in neurons using *alpha-tubulin:gal4* and more specifically in the Mauthner neuron, which regulates SLC behaviors, using the gal4 driver, *gffDMC130a* (**Supplemental Fig 2A-B**). Neither transgene restored normal levels of sensitivity or normal habituation in otherwise mutant animals, consistent with a CS-specific and non-neuronal role for *cdh16* in the regulation of acoustic startle thresholds.

**Figure 5.**
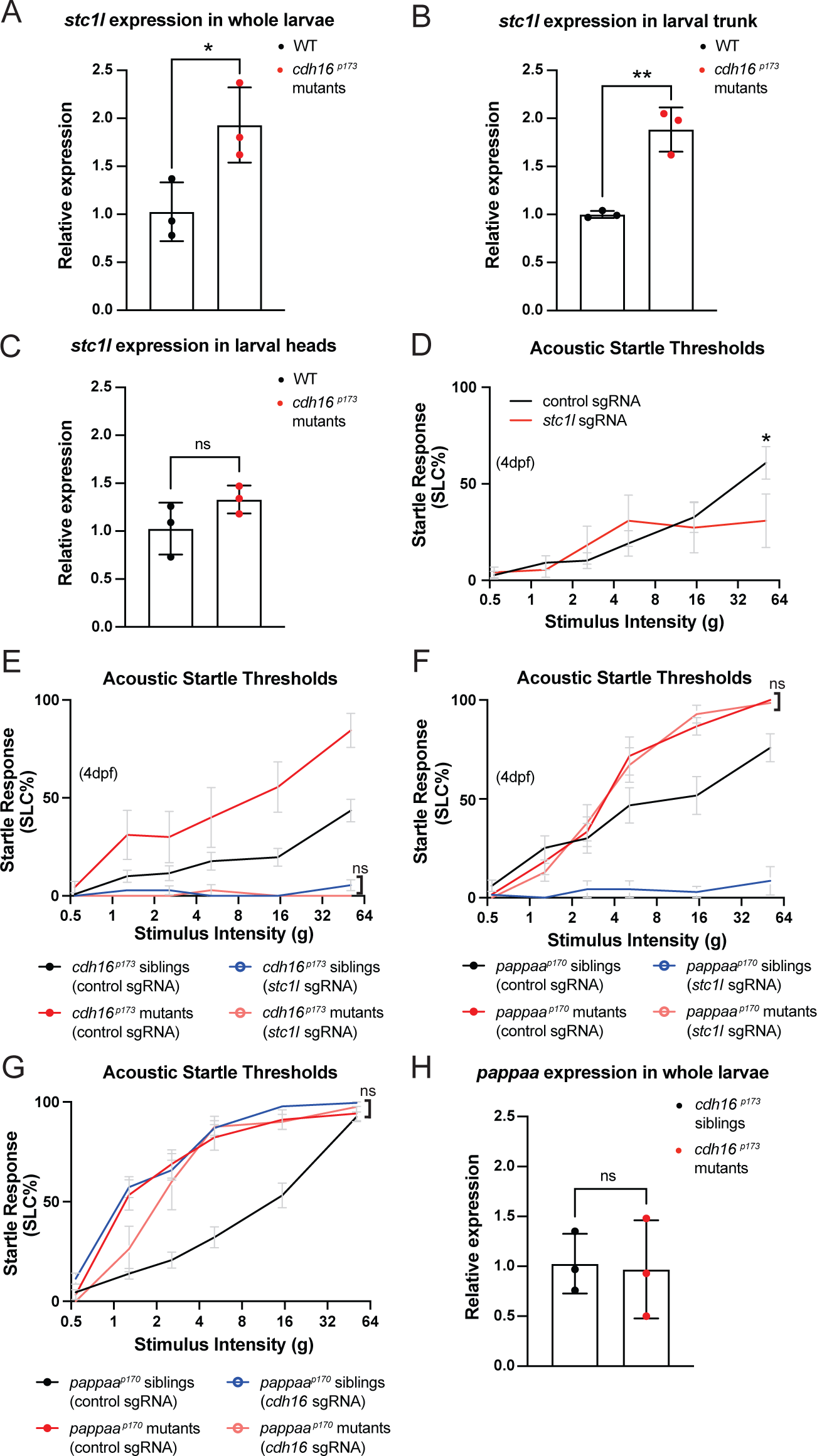
Genetic epistasis experiments reveal interactions between *cdh16*, *stc1l*, and *pappaa*. **(A-C)** RT-qPCR analysis of *stc1l* expression in *cdh16^p173^* mutants. **A)** Expression of *stc1l* mRNA is significantly increased in *cdh16* mutants compared to siblings. n=3 biological replicates per condition, *p=0.03 unpaired t-test. Error bars represent SD. **(B-C)** The increase in *stc1l* expression in *cdh16* mutants is observed specifically in trunk tissue, which includes the distal pronephros and CS **(B)** (n=3 biological replicates per condition, **p=0.003, unpaired t-test) and not the head **(C)** (n=3 biological replicates per condition, ns indicates p=0.16, unpaired t-test). Error bars represent SD. **D)** *stc1l* crispants (n=18) have a decreased response to acoustic stimuli compared to control guide-injected siblings (n=18) *p=0.0475, two-way ANOVA with Šídák’s multiple comparisons test. Error bars represent SEM. **E)** Genetic epistasis to examine the relationship between *stc1l* and *cdh16* in the context of acoustic startle thresholds*. stc1l* mutations suppress the *cdh16* mutant phenotype. *cdh16* mutants injected with *stc1l* guides (n=10) are not more responsive than siblings injected with *stc1l* guides alone (n=44) p>0.9 for all stimulus intensities, two-way ANOVA with Tukey’s multiple comparisons test. Error bars represent SEM. **F)** *pappaa* mutations suppress the *stc1l* crispant phenotype*. stc1l* guide-injected *pappaa* mutant larvae (n=14) are no more hyposensitive than control guide injected *pappaa* mutants (n=12) p>0.9 for all stimulus intensities, two-way ANOVA with Tukey’s multiple comparisons test. Error bars represent SEM. **G)** Loss-of-function mutations in *cdh16* and *pappaa* do not cause additive hypersensitivity phenotypes. *pappaa* mutants injected with *cdh16* guides (n=8) are no more hypersensitive than control guide injected *pappaa* mutants (n=18), p>0.8 at all intensities except for 1.3g, where p=0.0389, and control-guide injected are more sensitive than *cdh16* guide-injected *pappaa* mutants, two-way ANOVA with Tukey’s multiple comparisons test. **H)** *pappaa* mRNA levels are not altered in *cdh16* mutants (n=3 biological replicates per condition, p=0.87, unpaired t-test). Error bars represent SD.

Then, we predicted that since hypersensitive *cdh16* mutants overexpress *stc1l*, loss-of-function in *stc1l* would lead to hyposensitivity to acoustic stimuli. To test this, we performed F_0_ CRISPR mutagenesis experiments, injecting otherwise wild type embryos at the 1-cell stage with Cas9 together with either 3 control guides^45^ or together with 3 guides that we designed against *stc1l*. We found that loss of function in *stc1l* leads to severe pericardial edema, which becomes apparent by 5dpf as previously described^20^. Therefore, we tested behavior in larvae injected with *stc1l* guides (*stc1l* crispants) at 4dpf, before severe pericardial edema develops. At 4dpf, wild type zebrafish larvae are less responsive to acoustic stimuli, but as predicted, we found that *stc1l* crispants were even less responsive to acoustic stimuli than their control guide injected siblings (**Fig 5D**). To test whether *stc1l* overexpression in *cdh16* mutants is the cause of the hypersensitivity phenotype, we then performed the same CRISPR-Cas9 F_0_ mutagenesis in *cdh16* mutants and siblings. Consistent with our model, loss-of-function in *stc1l* in *cdh16* mutants resulted in hypo-responsiveness to acoustic stimuli (**Fig 5E**).

Previous work shows that Stanniocalcin limits Ca^2+^ uptake by inhibiting Papp-aa^20,42,43^. Therefore, we predicted that *pappaa* loss-of-function would suppress the hyposensitive phenotype observed in *stc1l* crispants, and that loss-of-function of both genes would resemble single mutants for *pappaa*. Indeed, we found that *pappaa* mutants injected with *stc1l* guides were hypersensitive, showing no difference relative to control-guide injected *pappaa* mutants (**Fig 5F**). If the function of *cdh16* is to release *pappaa* from inhibition by inhibiting *stc1l,* then animals carrying loss-of-function mutations in both *cdh16* and *pappaa* should be no more hypersensitive to acoustic stimuli than single mutants for either gene. Indeed, our crispant experiments are consistent with this model, as *pappaa* mutants injected with guides against *cdh16* were no more hypersensitive than *pappaa* mutants injected with control guides (**Fig 5G**). Finally, we set out to understand the mechanism through which *stc1l* regulates *pappaa* in the context of acoustic startle response thresholds. Stanniocalcin can both downregulate the expression of *pappaa*^20^ and separately inhibits its enzymatic function^42,43^. Consistent with a model in which increased expression of *stc1l* leads to inhibition of the enzymatic function of *pappaa*, we found that *pappaa* RNA expression levels were not changed in our *cdh16* mutants (**Fig 5H).**

### The corpuscles of Stannius and Ca^2+^ homeostasis are crucial regulators of acoustic sensory thresholds

Thus far, our data are consistent with a model in which *cdh16* and *pappaa* regulate Ca^2+^ homeostasis to promote acoustic startle thresholds and habituation. Importantly, in addition to its expression in Ca^2+^-regulatory ionocytes, *pappaa* is expressed in the supporting cells surrounding neuromasts, as well as in the retina and brain, including in the acoustic startle circuit^11,22,44,46,47^. However, it is not yet known whether *pappaa* expression in the brain or potentially in the ionocytes regulates sensory thresholds. First, to test whether *cdh16* mutants are hypocalcemic, we performed a colorimetric assay for whole-body Ca^2+^ content. Consistent with a model in which loss of *cdh16* leads to excessive *stc1l*, which downregulates *pappaa* and ionocyte proliferation and function to ultimately impair Ca^2+^ uptake, we found that *cdh16* mutants are hypocalcemic relative to their siblings (**Fig 6A**). Next, prior work has demonstrated that zebrafish raised in high-calcium media are hyposensitive to acoustic stimuli^48^. We wondered whether low-calcium media might produce larvae that are hypersensitive to acoustic stimuli. Indeed, we found that acute exposure to low-calcium media (0.001mM) resulted in acoustic hypersensitivity (**Fig 6B**) and animals exposed to this treatment trended towards a failure to habituate (**Fig 6C**). We note that exposure to 0.02mM Ca^2+^ caused the opposite phenotype: animals were hyposensitive and trended toward improved habituation. We speculate that this unexpected result may reflect engagement of compensatory mechanisms that drive animals toward hyposensitivity and note that wild type zebrafish larvae show a remarkable ability to cope with low environmental Ca^2+^ in terms of maintaining bone-mineralization^49^. Nonetheless, the specific mechanism underlying the complexity in the behavioral response to lowered Ca^2+^ remains unexplained. We additionally examined visually evoked behaviors (**Fig 6D, Supplemental Fig 3A-B**). Like *pappaa* mutants, animals exposed to low-calcium media (0.001mM) show reduced responsiveness to dark-flash stimuli, consistent with low calcium in *pappaa* mutants as an important driver of both phenotypes (**Fig 6D**). These data highlight that low Ca^2+^ and loss of *pappaa* both cause reduced escape responses to whole-field loss of illumination and increased responsiveness to acoustic stimuli.

**Figure 6.**
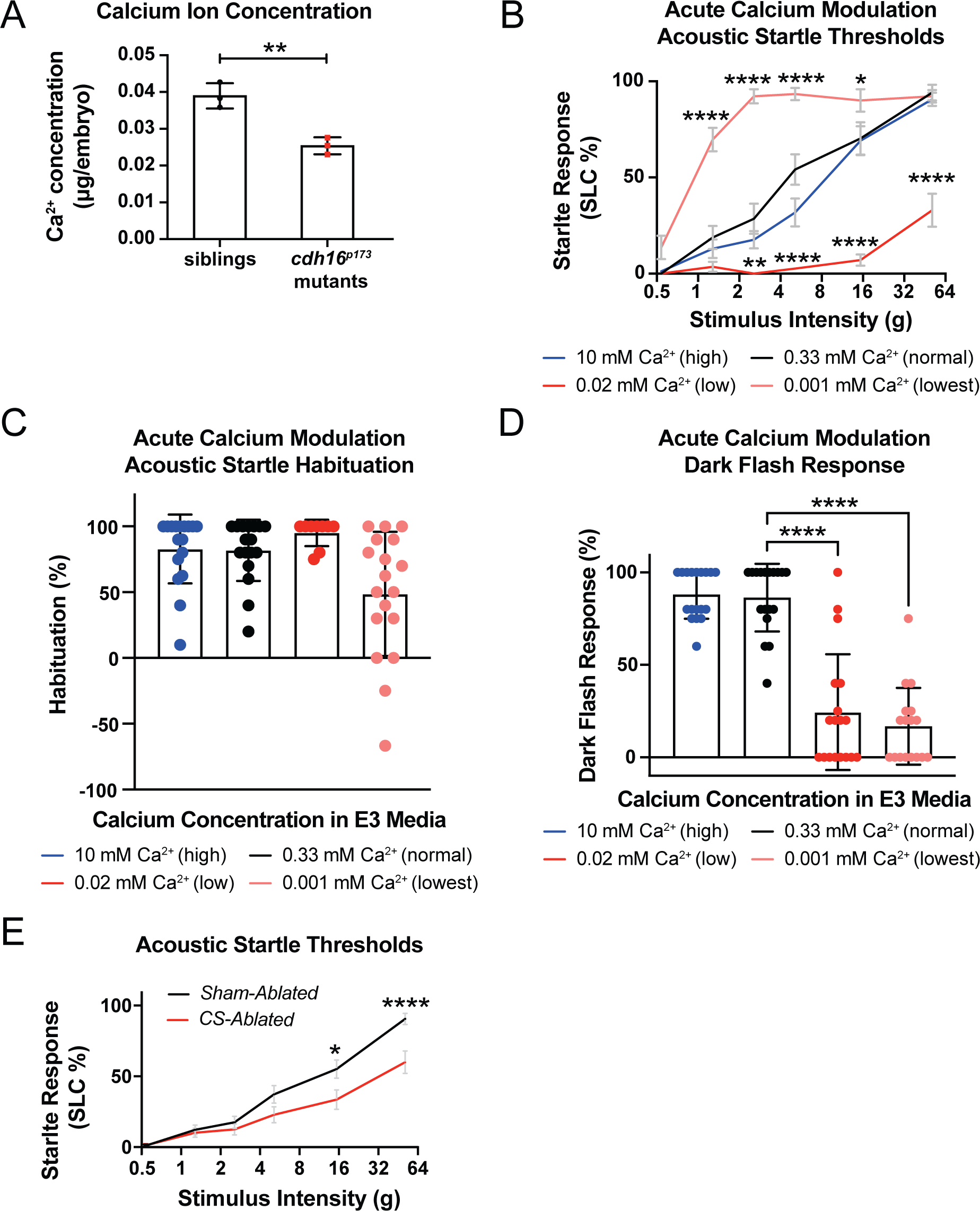
Calcium (Ca^2+^) homeostasis is critical for thresholding sensory-evoked behaviors. **A)** *cdh16* mutants have decreased whole-body Ca^2+^ compared to siblings (n=3 biological replicates per condition, **p=0.0048, unpaired t-test). Error bars represent SD. **B)** Larvae exposed to the lowest calcium media (0.001mM Ca^2+^) four hours before behavior testing have altered responses to acoustic stimuli. This condition increased sensitivity to acoustic stimuli at almost every stimulus intensity in larvae (n=18) compared to larvae (n=17) exposed to normal levels of Ca^2+^ (0.33mM); ****p<0.0001, *p=0.04. Larvae exposed to an intermediate-low level of Ca^2+^ (0.02mM, n=17) conversely, have reduced responses to acoustic stimuli relative to normal calcium (0.33mM) (n=17) **p=0.001, ****p<0.0001, two-way ANOVA with Dunnett’s multiple comparison’s test. Error bars represent SEM. **C)** Larvae in the lowest concentration of calcium trended towards a failure to habituate to acoustic stimuli (n=18) relative to larvae exposed to normal levels of Ca^2+^ (0.33mM, n=17) p=0.09, Kruskal-Wallis test with Dunn’s multiple comparisons test. Error bars represent SD. **D**) As is observed in *pappaa* mutant larvae, WT larvae exposed to low calcium (0.001mM, n=17) show decreased responding to dark flashes relative to larvae exposed to normal levels of Ca^2+^ (n=18) ****p<0.0001, Kruskal-Wallis test with Dunn’s multiple comparisons test. Error bars represent SD. **E)** Laser-ablation of the calcium-regulating corpuscles of Stannius (CS) causes decreased sensitivity to acoustic stimuli (n=20), compared to sham ablated siblings (n=20) *p=0.012, ****p<0.0001, two-way ANOVA with Šídák’s multiple comparisons test. Error bars indicate SEM.

Finally, our data suggest that *cdh16* regulates sensory thresholds through its function in the CS. To test this, we used a 532nm pulse laser to ablate the CS in otherwise wild type animals expressing *her6:mCherry*^50^, a transgene that labels the CS at 3-4dpf. Based on the overexpression of *stc1l* in the CS of *cdh16* mutants, and the suppression of hypersensitivity in *cdh16^p173^; stc1l* crispants, we predicted that ablation of the CS would result in hyposensitivity similar to that observed in *stc1l* crispant animals. Importantly, CS-ablated animals largely did not display pericardial edema at 5dpf (**Supplemental Fig 3C**). Those with pericardial edema were excluded from analysis. Consistent with a function for *cdh16* in the CS, we found that compared to their sham-ablated counterparts, CS-abated wild type animals were hyposensitive to acoustic stimuli (**Fig 6E, Supplemental Fig 3D-G**).

## Discussion

Taken together, our results highlight the corpuscle of Stannius as a brain non-autonomous endocrine regulator of sensory thresholds. Moreover, our results identify Cadherin 16 as an important regulator of endocrine function and highlight calcium homeostasis as critical for sensory gating *in vivo*. Based on our data, we propose that without *cdh16*, Stanniocalcin 1L is overexpressed, PAPP-AA function in the proliferation of ionocytes and/or expression of the calcium channel *trpv6* is suppressed, and insufficient Ca^2+^ is taken up from the environment. The ultimate consequence is that zebrafish larvae are hypocalcemic, leading to hypersensitivity to acoustic stimuli and in the case of *pappaa* loss-of-function, insufficient responding to whole-field loss of illumination (dark flash response) (**Fig. 7A-B**).

**Figure 7.**
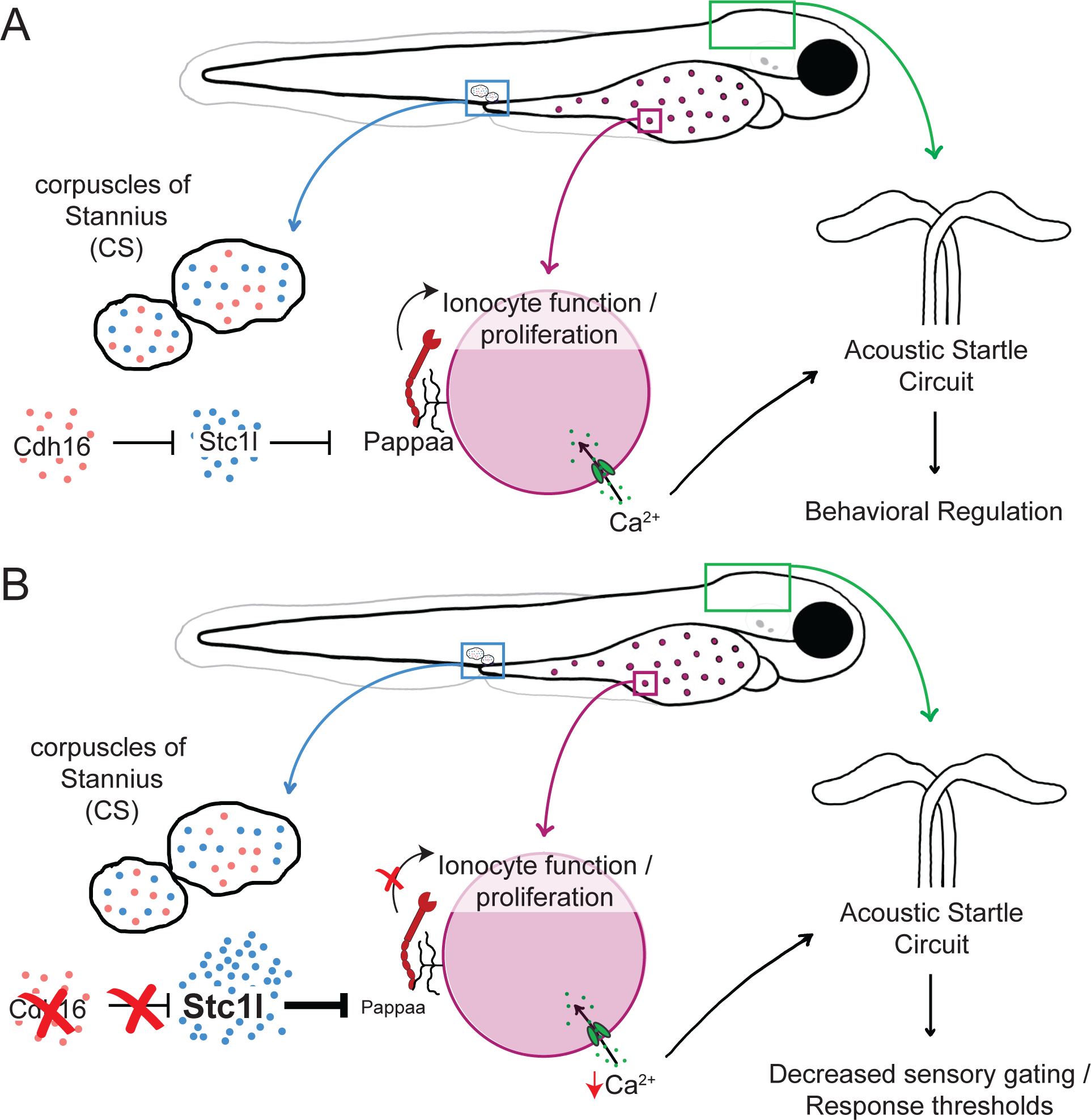
Proposed Model. **A)** In wild type animals, *cdh16* suppresses *stc1l* expression in the corpuscles of Stannius. This limits the ability of *stc1l* to suppress the function of PAPP-AA (we propose at the level of ionocytes), allowing for some proliferation and function of ionocytes. As a result, Ca^2+^ is taken up from the environment and normal acoustic startle thresholds are maintained. **B)** In *cdh16* mutant animals, suppression of *stc1l* expression is relieved and *stc1l* is overexpressed. This results in hyperinhibition of PAPP-AA. As a result, Ca^2+^ uptake is severely limited, animals are hypocalcemic, and acoustic response thresholds are lowered.

We have not yet established whether overexpressed Stanniocalcin1L suppresses *pappaa* function at the level of ionocytes to regulate behavior. It is possible that this is the key locus for their interaction, but *pappaa* is also expressed in supporting cells of the lateral line neuromasts and in the retina^44,46,47^. Therefore, it’s possible that *stc1l* impacts *pappaa* function within one or a combination of these structures. No matter where this key pathway functions, these data provide a parallel with human patient data indicating that hypocalcemia is associated with disruptions in auditory gating^24,25,51^.

We found that *cdh16* regulates acoustic thresholds and habituation after the development of the acoustic startle circuit and after the CS is established. Restoration of *cdh16* expression at 4 and 5dpf reverts behavioral deficits such that responding is normal at 6dpf. Similarly, *pappaa* function is sufficient later in development. Restoration of PI3K signaling downstream of *pappaa* at 5dpf restores habituation^11^. Low-calcium exposure also causes hypersensitivity independent of early development: acoustic hypersensitivity is apparent after only 4 hours in low Ca^2+^ media in 5dpf fish. Similarly, in patients with hypocalcemia, psychotic symptoms are locked to periods of calcium dysregulation, and normalization of Ca^2+^ levels can normalize symptoms^24,25^. These data extend previous findings that developmental exposure to Cadmium (an inhibitor of Ca^2+^ channel function) impacts sensory thresholds^52^, indicating that even acute disruptions in Ca^2+^ homeostasis can impact behavior.

We do not yet know precisely how hypocalcemia impacts activity within the neuronal circuits responsible for gating sensory stimuli^53^. In hippocampal slices, low Ca^2+^ exposure results in an increase in spontaneous neuronal activity^54^. This effect may be partially explained by a somewhat depolarized resting membrane potential mediated by depolarizing currents through sodium leak channels (NALCN) under conditions of hypocalcemia^55^. In this model, Ca^2+^ is detected by the calcium sensing receptor CaSR, which suppresses current through NALCN^55^. Under conditions of low Ca^2+^, NALCN currents are dis-inhibited and neurons are somewhat depolarized. Signaling through CaSR separately regulates firing frequency through regulation of Calcium-Activated Potassium Channels^56^.

Interestingly, loss-of-function mutations in the Calcium sensing receptor, *casr* were also uncovered in the forward genetic screen for regulators of acoustic startle response gating^12^. Like *cdh16*, CaSR regulates whole-body calcium levels, but in humans, patients with inactivating mutations in CaSR are hypercalcemic^57^ (in contrast to *cdh16* mutants, which we showed are hypocalcemic). Mirroring their opposing impacts on Ca^2+^ homeostasis, CaSR and Cdh16 have somewhat opposing impacts on behavior. While *cdh16* mutants are hypersensitive to acoustic stimuli and perform more short-latency startles, *casr* mutants perform fewer short-latency startles, instead responding to acoustic stimuli by primarily performing a distinct behavior, the long-latency C-bend, which wild type zebrafish larvae ordinarily perform in response to lower-intensity stimuli^12^. However, the role of CaSR is likely more complex. In addition to regulating serum Ca^2+^, CaSR functions in neurons to regulate acoustic startle response gating^16^. Restoration of CaSR function in otherwise *casr* mutant animals in a small population of hindbrain neurons that project in the vicinity of the Mauthner cell restores normal startle responsiveness^16^. Presumably, these rescued animals remain hypercalcemic, but rescue of CaSR signaling within this particular population is sufficient to normalize behavior. How and if the Cdh16, Stc1l, Papp-aa pathway interacts with CaSR signaling in the brain is not yet known, though we note that Papp-aa is expressed in multiple neuronal populations within the acoustic startle circuit^11,22^ and could interact with CaSR there.

Additional support for a link between the *pappaa* and *casr* pathways is provided by our recent work finding similar whole-brain activity patterns and drug response profiles for animals carrying loss-of-function mutations in *pappaa* and *ap2s1*^17^, which genetically interacts with *casr*^12^. Like *pappaa* and *cdh16*, *ap2s1* mutants are hypersensitive and fail to habituate to acoustic stimuli^11,58^, and mutations in *ap2s1* significantly suppress the CaSR phenotype^12^. In light of our new data connecting *pappaa* to *cdh16* and Ca^2+^ homeostasis, and *ap2s1*’s genetic interaction with the calcium-regulatory CaSR, we now propose that the commonalities between the *ap2s1* and *pappaa* whole-brain activity patterns may reflect common dysregulation of Ca^2+^.

*cdh16* and *pappaa* mutants, as well as wild type animals exposed to low Ca^2+^, show acoustic sensory gating deficits. Conversely, only low Ca^2+^-exposed fish and *pappaa* mutants exhibit deficits in the visually evoked O-bend response. Interestingly, in addition to its expression in ionocytes and neuromast support cells, *pappaa* is expressed in the retina, where mutants show disrupted development of synapses between photoreceptor cones and OFF bipolar cells^44^. *pappaa* mutants also have a thinner outer plexiform layer (the layer where cones make synaptic contacts with bipolar cells)^44^. Notably, in mice and zebrafish, mutations in *cacna1fa,* which encodes a Ca^2+^ channel essential for maintaining resting Ca^2+^ currents in photoreceptors, are also associated with visual defects and thinning of the outer plexiform later. Mutations in *pde6c*, which regulates Ca^2+^ channels in cones, are similarly associated with both visual defects and defects in the outer plexiform layer^59–61^. Finally, acute exposures of dissected mouse retinae to calcium chelators results in disassembly of presynaptic terminals in photoreceptors^62^, and acute inhibition of Ca^2+^ channels results in synaptic deficits in the zebrafish retina^59^. These data, together with our finding that low Ca^2+^ and loss of *pappaa* have the same effects on the response to dark flash, lead us to propose that disruptions in Ca^2+^ homeostasis may be responsible for the *pappaa* visual and acoustic phenotypes.

Finally, we still do not know how loss of *cdh16* leads to a rise in Stanniocalcin 1L expression. Cadherin 16 is an atypical cadherin within the 7-Domain family of cadherins and characterized by a short intracellular domain lacking binding sites for catenins. Therefore, although Cadherin 16 can function as an adhesion protein^32^, the intracellular mechanism underlying Cadherin 16 regulation of downstream *stc1l* expression is not yet known. Recent work shows that mutations in *sox10* increase the number of *stc1l*-positive cells in the CS, consistent with a possible role in regulating the proliferation of CS cells^49^. A similar mechanism might be at work in *cdh16* mutants. Perhaps *stc1l* is increased because more CS cells are present to produce *stc1l*. Whether *cdh16* directly regulates *stc1l* gene expression, or whether it suppresses *stc1l* indirectly via a primary effect on CS proliferation is not yet known. These questions are relevant to our understanding of sensory gating and the development of the CS, but also for cancer biology, as *cdh16* is downregulated in thyroid carcinomas^63^ and limits thyroid carcinoma cell proliferation^64^.

Taken together, our studies support a model in which Cdh16 suppresses Stc1l secretion from the CS, a role that it continues to play throughout larval development rather than during the specification or assembly of the CS. Stc1l then suppresses Papp-aa function, ultimately promoting hypocalcemia and responsiveness to acoustic stimuli. This work highlights a previously unappreciated role for Ca^2+^ homeostasis in the regulation of acoustic response thresholding and identifies a new brain non-autonomous pathway for the regulation of behavior.

## Materials and Methods

### Ethics statement

All procedures were approved by the University of Colorado Anschutz Medical Campus School of Medicine Institutional Animal Care and Use Committee (IACUC).

### Experimental Model and Subject Details

Zebrafish larvae were obtained from pairwise or group crosses of adult zebrafish carrying mutations or transgenes of interest on the TLF (WT) background. Larvae were raised at 28.5°C in E3 media and sorted for normal development.

The *p173* allele of *cdh16* and the *p170* allele of *pappaa* were recovered from a forward genetic screen^11^. Mutants were genotyped using proprietary allele specific primer sequences (LGC Genomics) and the KASP assay method, which utilizes FRET to distinguish between alleles. For genotyping of *p173* in the context of *Tg[hsp70:cdh16-p2a-mkate],* CAPS primers 107 and 108 were used in combination with MseI (see **Table 1**).

The *co79* mutant allele was generated using CRISPR-Cas9 mutagenesis. sgRNA 622 (**Table 1**) was designed using ChopChop^65^. The sgRNA was purchased from IDT and reconstituted to 200uM using the IDT-provided duplex buffer. sgRNA was combined with tracrRNA, also purchased from IDT, to form a 50uM duplex by heating at 95°C in a thermocycler for 5 minutes, followed by cooling to RT for 10 minutes. Injection mixes were prepared by mixing 1uL of 50uM duplex together with 1uL Cas9 protein (5mg/mL) obtained from PNA Bio and 1uL phenol red. *cdh16^co79^* mutations were genotyped by PCR with primers 657 and 658 (**Table 1**).

Transgenic animals carrying *Tg[hsp70:cdh16-p2a-mkate]* (*co113*) were generated by cloning the *cdh16* cDNA from total zebrafish RNA at 5dpf into *pME-cdh16-p2a-mKate*. Gateway cloning was used to recombine *pME-cdh16-p2a-mKate* into a pDest vector containing the *hsp70* promoter and I-sceI restriction sites, generating *hsp70-cdh16-p2a-mKate*. I-sceI transgenesis was performed as previously described^66^ by injecting I-sceI and the *hsp70-cdh16-p2a-mKate* plasmid into 1-cell stage TLF embryos. G_0_ injected larvae were raised, outcrossed, and heat-shocked at 37°C in a thermocycler for 45 minutes to identify carriers. Larvae expressing the transgene were identified by screening for mKate using a fluorescent stereomicroscope (Leica M205FCA). For behavior experiments, animals were pre-screened for fluorescence and genotyped post-hoc using primers 107 and 108 (**Table 1**).

Transgenic animals carrying the *gal4* driver *Tg[gffDMC130a]* were provided by the lab of Dr. Koichi Kawakami^67^. Transgenic animals carrying *Tg[alpha-tubulin:gal4]* were provided by the lab of Dr. Philippe Mourrain^68^. Animals carrying *Tg[her6:mCherry]*^50^ were provided by the lab of Dr. James Nichols and outcrossed to TLF for ablation experiments.

To generate conceptual translations of each allele, SMART domain-prediction software was used^69^. SMART identified Cadherin repeats 1-6 based on the full-length protein sequence. Cadherin repeat 7 was not originally identified, however SMART identified a 7^th^ cadherin repeat when the final portion of the extracellular domain was searched alone.

*Tg[UAS:cdh16-EGFP]* was made by injecting the *UAS-cdh16-EGFP* plasmid along with Tol1 transposase into *cdh16^p173^*+/−; *alpha-tubulin:gal4*+/0 embryos at the single-cell stage. Carriers were identified by screening for EGFP expression using a fluorescent stereomicroscope, raising positive offspring to adulthood, and outcrossing them to TLF. The *Tg[UAS:cdh16-EGFP]* fish were further outcrossed to *alpha-tubulin:gal4*+/− for examining neuronal *cdh16* expression. Larvae were pre-sorted for EGFP expression before behavior testing. After behavior testing, larvae were genotyped for *cdh16^p173^* using primers 107 and 108 (Table 1). To assess Mauthner cell-specific rescue, *Tg[UAS:cdh16-EGFP]; cdh16^p173+/−^* fish were crossed to *Tg[Gap43:Citrine]; Tg[*Gffdmc130a]; cdh16p173+/−. Larvae were genotyped after behavior testing for the rescue construct with primers 107 and 1002, and further genotyped for *cdh16^p173^* using primers 107 and 108 (**Table 1**).

### Behavior Testing

Before testing their response to acoustic and visual stimuli, larvae were acclimated to the behavior room inside an incubator kept at 28°C for 30 minutes. To measure acoustic startle thresholds, six increasingly intense acoustic stimuli were administered 5 times each, 40 seconds apart, after which acoustic startle response habituation was measured by providing 40 stimuli with a 3-second interstimulus interval (ISI). Visual motor responses were measured by first dark-acclimating larval zebrafish inside the behavior arena. Next, the lights were turned on for a 7-minute period to assess initial visual motor reactivity in response to light. Then, the lights were turned off for 7 minutes to assess the initial visual motor response to darkness. Light flash reactivity was examined by first dark-acclimating larval zebrafish inside the behavior arena. Next, larvae were exposed to ten pulses of light with a one second duration, 30 seconds apart. To assess dark flash reactivity, 6dpf larvae were acclimated to the light inside the behavior arena. Following this, the lights were extinguished 5 times in pulses lasting 1 second with a 1-minute ISI. To assess dark-flash habituation 60 additional dark flash stimuli were administered with a 10-second ISI. During these final stimuli, the camera recorded behavior during every other stimulus. For the above-described behavior assays, larvae were loaded onto a custom-made acrylic 6×6 well-plate attached to a mini-shaker (Brüler & Kjær, Model 4810), which was used to deliver the acoustic stimuli. A cover was placed over the rig for assays of visually evoked behaviors.

Behavior was recorded with a high-speed camera (FASTCAM Mini UX50 Type 160K-M-32G) placed above the plate and an internal LED light pointed at the behavior arena was used for light stimuli. Acoustic stimuli were calibrated using an accelerometer (PCB Piezotronics, Y355B03) and stimulus intensities are reported in g or acceleration due to gravity. To analyze behavior, video files were background-subtracted and then analyzed using FLOTE, Batchan^26^, and Microsoft Excel. Statistical analyses and graphing were performed using Graphpad Prism.

Larvae were tested for thermal behavior using a 96-well (square wells) plate loaded into a DanioVision observation chamber running EthoVision XT 11.5 software (observation chamber and software, Noldus, Leesburg, VA). The temperature in the observation chamber was set using a temperature control unit. Larvae were acclimated to the baseline temperature of 28.5°C for 30 minutes, after which their total distance moved was recorded for 2 minutes. The temperature was then raised to 33.5°C, and fish were recorded again for 2 minutes. All behavioral assays were performed at 5dpf, except for our dark flash assay which was performed at 6dpf.

### Heat-shock induced *cdh16* rescue

To induce expression of *hsp70-cdh16-p2a-mKate*, zebrafish embryos or larvae were placed in a 96-well plate at a density of no more than 5 larvae per well. The plate was heated to 37°C for 45 minutes using a thermocycler. Larvae were then recovered to petri dishes for at least 5 hours before behavior testing.

### Crispant (F_0_) Mutagenesis and Behavior Analysis

sgRNAs targeting *cdh16* (622, 623, 867) and *stc1l* (942, 943, 944) were designed using ChopChop^65^. Scrambled sgRNAs (759, 760, and 761) were used as controls and were designed by IDT as previously described^45^. The sgRNAs were purchased from IDT and reconstituted to 200uM stocks using the IDT-provided duplex buffer. sgRNAs were then combined individually with tracrRNA, also purchased from IDT, to form a 61uM duplex by heating at 95°C in a thermocycler for 5 minutes, followed by cooling to RT for 10 minutes. Injection mixes were prepared by mixing 1uL of duplex together with 1uL Cas9 nuclease V3 (10ug/uL; IDT Cat #1081059). 1nl of injection mix was injected in the yolk at the single cell stage, before the cell inflates.

The mutation rate in crispants was assessed by PCR using primers flanking the sgRNA target sequences to detect indels and large deletions (see Table 3). Following behavioral analysis, we genotyped larvae injected with gene-specific sgRNAs and larvae injected with control sgRNAs to confirm guide efficiency.

### Hybridization Chain Reaction (HCR) FISH staining

HCR probes, hairpins, and buffers were purchased from Molecular Instruments. Staining was performed using the manufacturer’s protocol: “HCR RNA-FISH protocol for whole-mount zebrafish embryos and larvae (*Danio rerio*)” with the following modifications: we did not apply PTU to inhibit melanogenesis, we used 30 larvae per Eppendorf tube, and we used 8uL of 1uM *cdh16* probe solution instead of 2ul as suggested in the protocol. Animals were mounted laterally in 1.5% low-melt agarose in PBS and imaged using a 63x objective on a 3i Marianas Spinning Disk Confocal Microscope.

### Calcium manipulations

To create calcium-supplemented media, we first created a stock solution of 60x E3 embryo media without calcium: 300 mM NaCl, 10.2 mM KCl, and 19.8mM MgSO_4_·7H_2_O. A separate stock solution of 60x CaCl_2_·2H_2_O (Sigma CAS#:10035-04-8) was also made. Calcium concentrations of 10mM Ca^2+^, 0.33mM (Normal), 0.02mM, and 0.001mM were generated by mixing 60x E3 and 60x CaCl_2_ in the appropriate ratios. At 5dpf, larvae were rinsed three times out of E3 media containing normal calcium (0.33mM Ca^2+^), and into one of the four different calcium-supplemented media concentrations four hours before performing behavior and then tested in those same calcium concentrations.

### Corpuscle Ablations

*Tg*[*her6:mCherry]* embryos were screened for mCherry expression at 3dpf using a fluorescent stereomicroscope. 4dpf mCherry-positive larvae were live-mounted laterally in 1.5% low-melt agarose (Lonza Cat# 50101) in E3 embryo media on a 3.5 cm glass-bottom dish. The CS were identified and then ablated using 532 nm pulse laser attached to a 3i Marianas spinning disk confocal microscope with a 63x objective. To ensure complete ablation, an average of 3 laser pulses were administered per corpuscle (laser pulses were delivered until the CS was eliminated). For sham ablations, a target region posterior to the kidney and yolk extension was located and ablated, after which the CS were re-located and confirmed to be undamaged. Ablated and sham-ablated larvae were then unmounted and placed in a 6cm petri dish with fresh E3 to recover for approximately 21 hours, after which they were behavior tested at 5dpf for acoustic startle thresholds and habituation.

### WGS and Molecular Cloning of *cdh16*

Molecular cloning of the *cdh16* allele was performed as previously described^11,14^. Pools of 50 behaviorally identified *p173* mutant larvae were collected and used to prepare genomic DNA (gDNA) libraries. gDNA was sequenced with 100-bp paired-end reads on the Illumina HiSeq 2000 platform, and homozygosity analysis was done using 463,379 SNP markers identified by sequencing gDNA from ENU-mutagenized TLF and WIK males as described previously^11^.

### Calcium Content Assays

Whole-body Ca^2+^ was quantified using a colorimetric assay kit (Abcam ab102505). 2dpf larvae were live tail-clipped and genotyped for the *cdh16^p173^* allele. At 4dpf, 10-15 larvae were pooled in 6 Eppendorf tubes: three WT biological replicates and three mutant biological replicates. The assay was then performed as previously described^49^.

### RT-qPCR

Larvae were dissected to remove the distal tip of the tail for genotyping. To generate cDNA from heads and trunks, larvae were dissected to isolate the head from the trunk at the base of the hindbrain. Tissue to be used for RT-qPCR was placed into RNAlater (Sigma Cat# R0901-100ML) and stored at 4°C. Following genotyping, whole larvae (**Fig 5A, 5H**), trunks (**Fig 5B**), or heads (**Fig 5C**) were pooled by genotype (homozygous *cdh16^p173^* mutants and homozygous wild type siblings, 10 larvae per pool, 3 biological replicates) and total RNA was extracted using Trizol/Chloroform followed by the RNeasy Plus Mini Kit (Qiagen Cat# 74143). cDNA pools were generated using SuperScript II Reverse Transcriptase (Invitrogen Cat# 11904-018). qPCR was performed with LUNA qPCR MasterMix (NEB Cat# M3003) on a QuantStudio 3 Real-Time PCR System (Fisher Cat# A28566) using qPCR primers (**Table 4**) designed for each target gene. Expression levels of target genes were normalized to *gapdh*.

### Quantification and Statistical Analysis

Statistical tests were performed in Graphpad PRISM 9 and 10. To determine normality for each data set, the D’Agostino & Pearson test was performed. In normally distributed data, an unpaired T test, one-way ANOVA, or two-way ANOVA was performed as needed. To account for multiple comparisons in two-way ANOVAs, the Šidák’s multiple comparisons test was performed when comparing means across one variable while the Tukey’s multiple comparisons test was used to compare means between all experimental groups. In datasets that are not normally distributed, a Mann-Whitney test was executed to compare two groups and a Kruskal-Wallis test was used to compare between greater than two groups.

### Whole Brain Morphometric Imaging and Analysis

6dpf larvae (n=60) from a *cdh16^p173^* heterozygote incross were acclimated to the behavior testing room for 30 minutes. Following acclimation, the larvae were placed in a cell strainer within a 6cm petri dish containing E3 for 30 minutes. Finally, spontaneous behavior was recorded for 16 minutes before the cell strainer was removed and placed into a 6 well dish containing 4% paraformaldehyde in PBT (PBS-Triton 0.25%) for 45 seconds to flash-fix the larvae. The cell strainer was then transferred to a solution of 4% paraformaldehyde in PBS, incubating at 4°C overnight. Larvae were moved from the cell strainer to a 1.5mL tube and washed with PBT for three, 5-minute washes. To increase the ratio of mutants to WT larvae included in the imaging experiment, tail clips were collected from each sample, lysed, and KASP genotyped for the *cdh16^p173^* mutation. Wild type and mutant larvae were pooled at a 1:1 ratio into a 1.5mL tube containing PBT and stained according to a previously developed immunohistochemistry protocol for MAP-mapping^37^ with procedural alterations^17^. Finally, samples were mounted onto a glass-bottom dish using 1.5% low-melt agarose made with PBS. Each larva was positioned with the dorsal portion of its brain facing the glass bottom of the dish. Whole-brain z-stacks were collected for each sample using an LSM780 microscope with a 20x objective and 2×1 tile scanning. Larvae were unmounted from the agarose and gDNA was prepared for KASP genotyping. Morphometric analysis of *cdh16^p173^* mutants was then performed as previously described^36,37^. Differences in whole brain morphology were examined by assessing the significant delta medians of mutants over WT.

### Genotyping Table 1

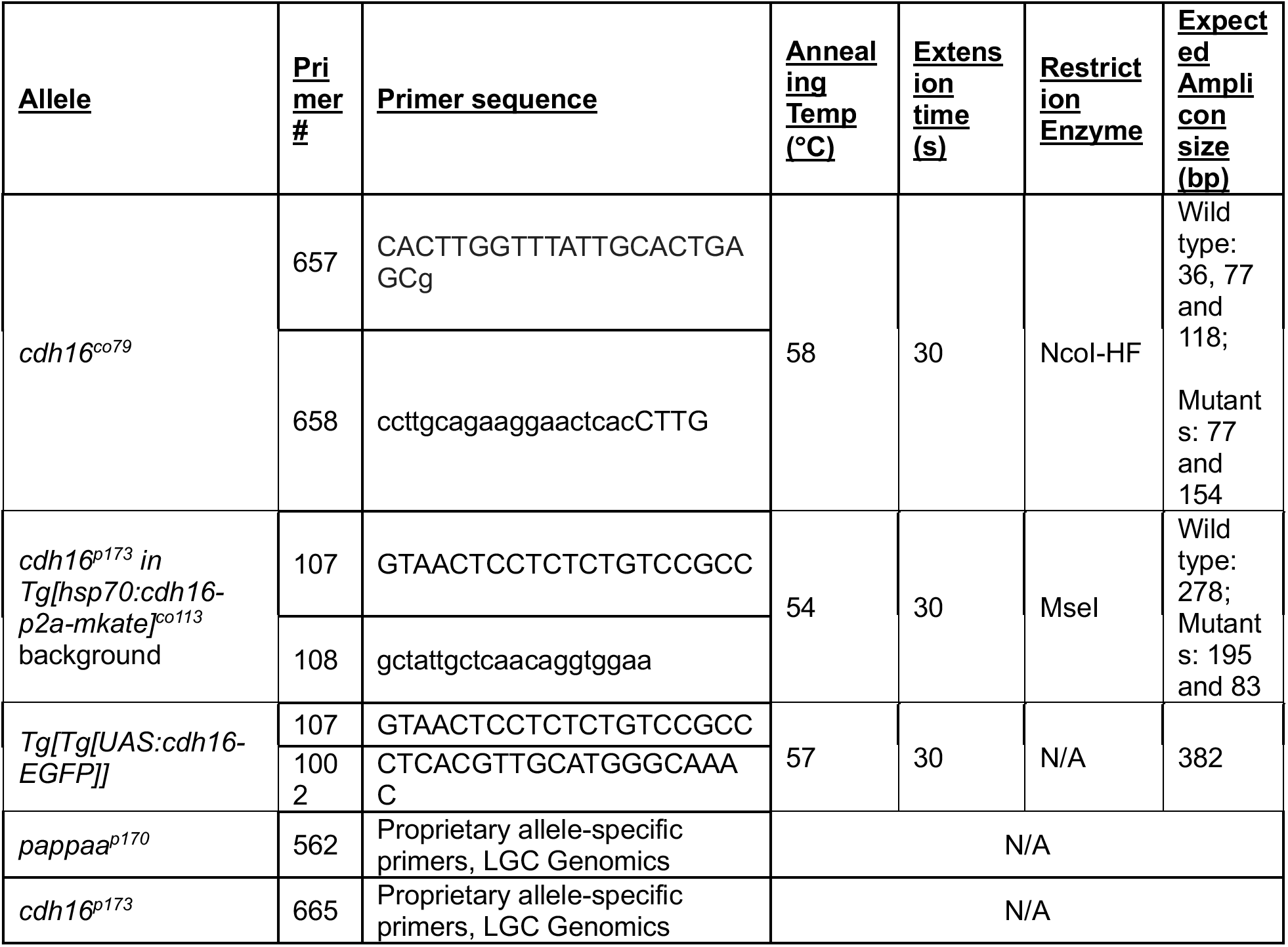

### Cloning Primers Table 2

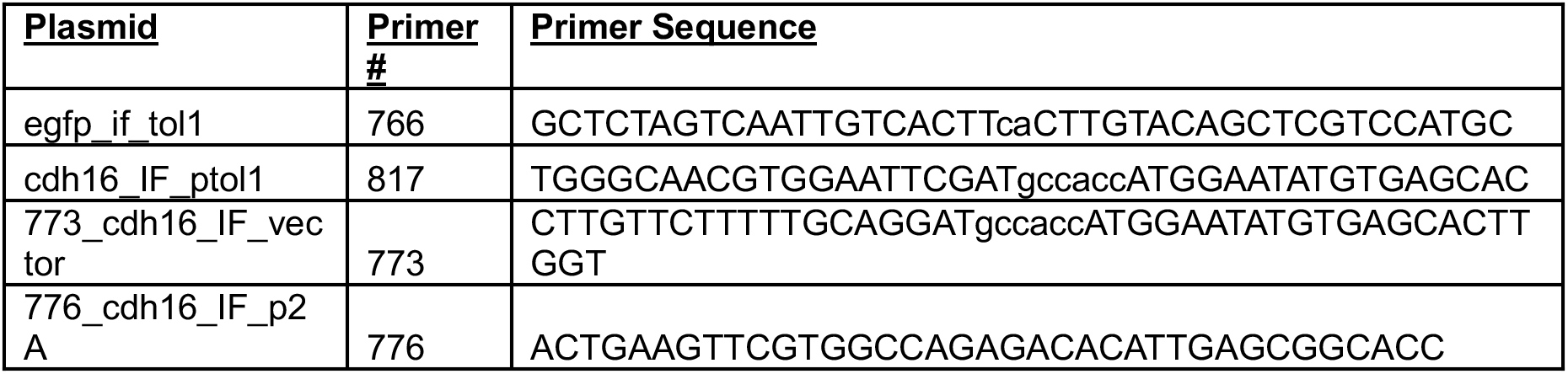

### Analyzing CRISPR Efficiency Table 3

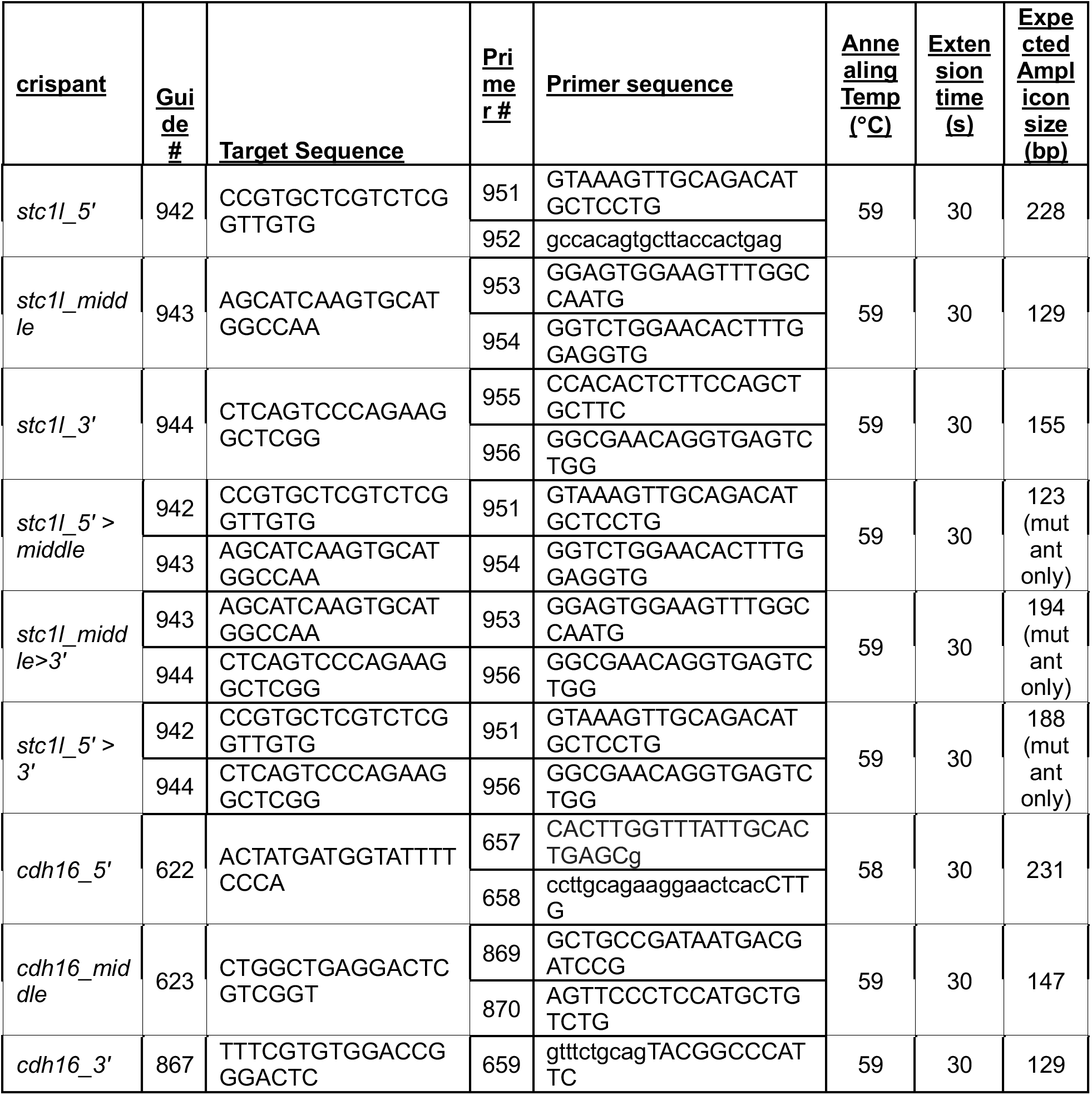
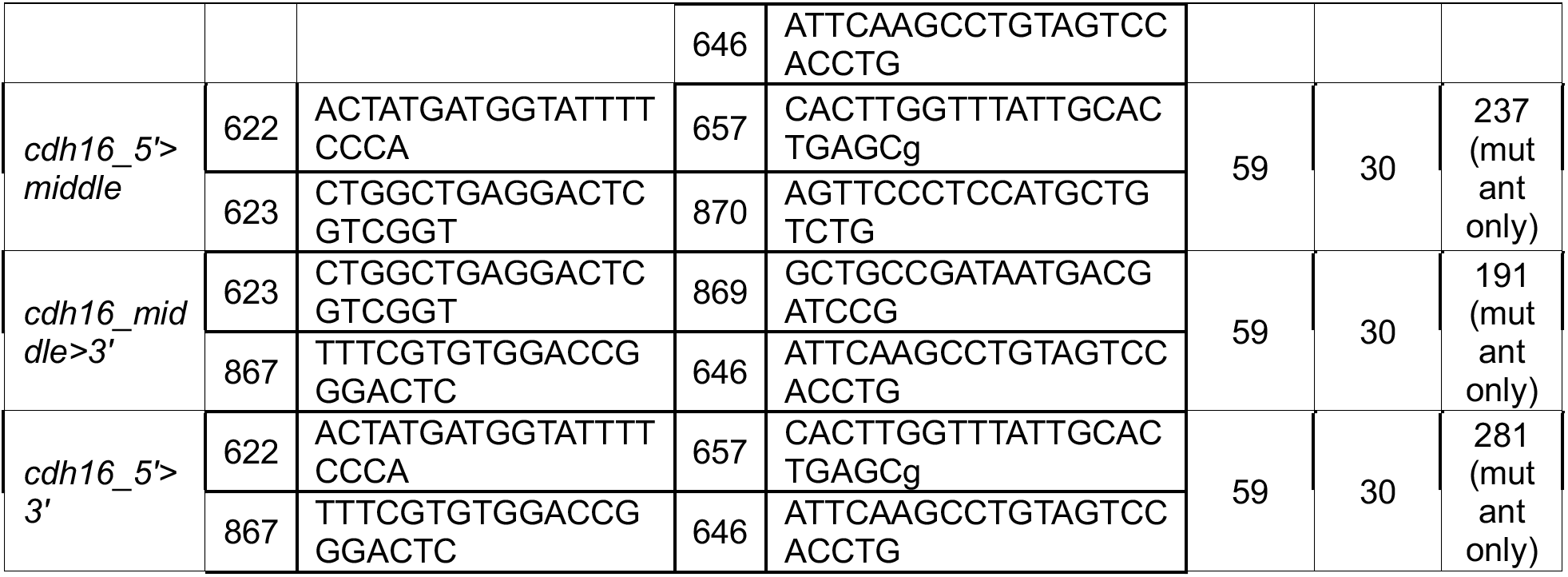

### qPCR Primers Table 4

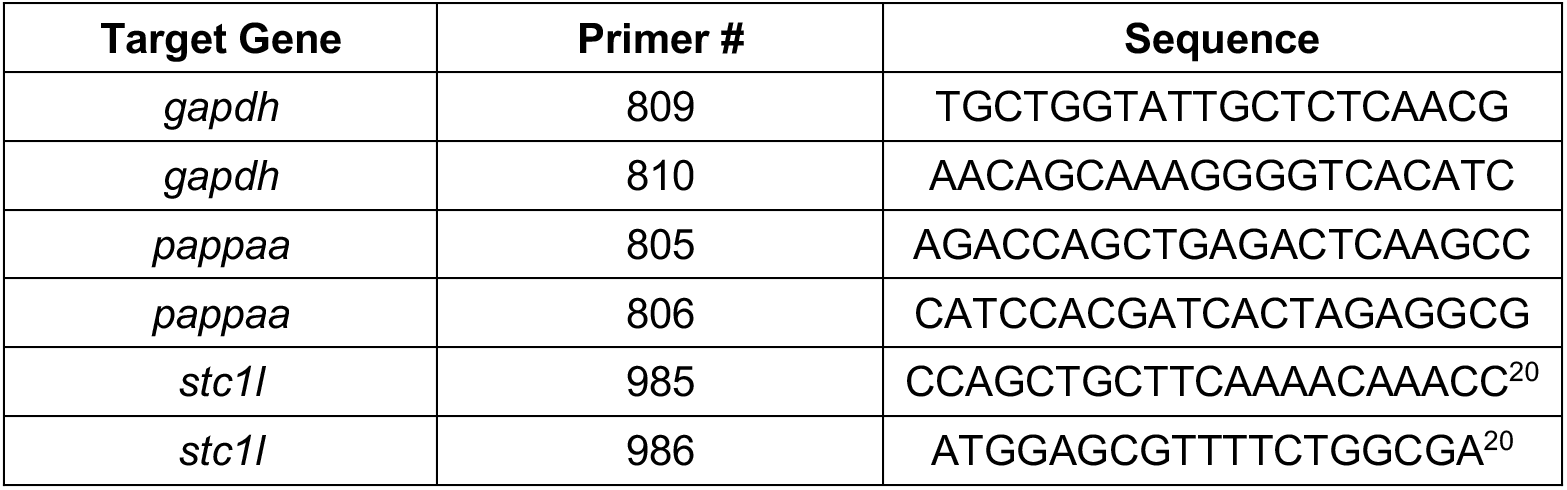

## Acknowledgements

We are grateful to Dr. Michael Granato, Dr. Marc Wolman, Dr. Roshan Jain, Dr. Kurt Marsden, Hannah Bell, Julianne Skinner, Katharina Hayer, and Dr. John Hogenesch for conducting the original ENU screen, and developing the whole genome sequencing analysis pipeline. We thank Dr. Hannah Shoenhard, Dr. Joy Meserve, and Dr. Caleb Doll for thoughtful comments on a draft of this manuscript. We are grateful to Dr. James Nichols, Colette Dolby, Abi Mumme-Monheit, and other members of the Nichols lab for sharing *her6:mcherry* transgenic zebrafish as well as technical assistance and suggestions. We thank Dr. Summer Thyme and Ari Ginsparg for assistance with morphometric analyses. We thank Dr. Emerald Butko for sharing the observation that *her6:mcherry* labels the pronephric duct. We thank the University of Colorado Anschutz Medical Campus Zebrafish Facility, the University of Colorado Anschutz Medical Campus NeuroTechnology Center’s Advanced Light Microscopy Core, and The University of Colorado Anschutz Medical Campus NeuroTechnology Center’s Animal Behavior & In Vivo Neurophysiology Core. This work was supported by funds from the Boettcher Foundation’s Webb-Waring Biomedical Research Awards program. Funding was also provided by the NIH / NINDS R00NS111736 (awarded to JCN), the Center for Pediatric Genomics at Cincinnati Children’s Hospital (awarded to LB), and the NIH T32 GDDR (awarded to SSS).

## Author Contributions

Conceptualization, SSS, and ZQM, JCN; Methodology, SSS, ZQM, LB, and JCN; Investigation, SSS, ZQM, NJS, SG, AS, LB, JCN; Resources, SSS, LB, JCN; Writing – Original Draft, SSS, ZQM, NJS, JCN; Writing – Review & Editing, all authors, Supervision, LB, JCN.

---

**Supplemental Figure 1.**
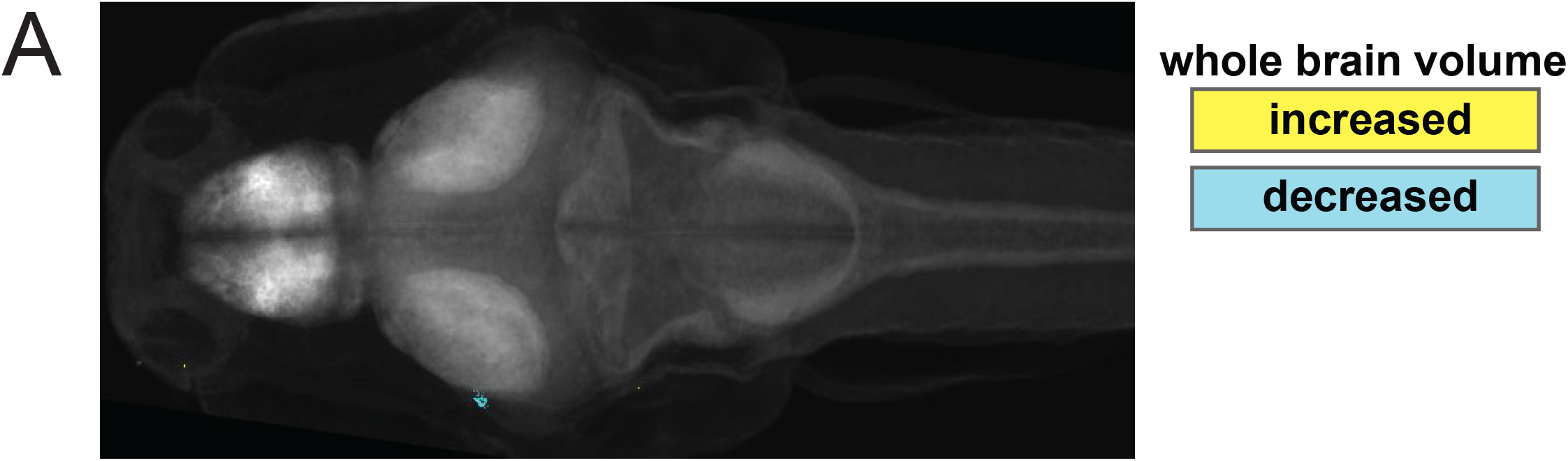
Whole-brain morphometric analysis reveals minimal changes to region-by-region brain volume. **A)** Summary of whole-brain morphometric data for 6dpf *cdh16^p173^* mutants (n=13) as compared to siblings (n=19). Region-by-region differences in volume are indicated in yellow (regions that are larger in mutants) or cyan (regions that are smaller in mutants). Image is a summed stack of the significant delta medians of mutants over wild types. Note there are no colored pixels within the brain, indicating no significant differences between mutants and siblings across the annotated brain regions.

**Supplemental Figure 2.**
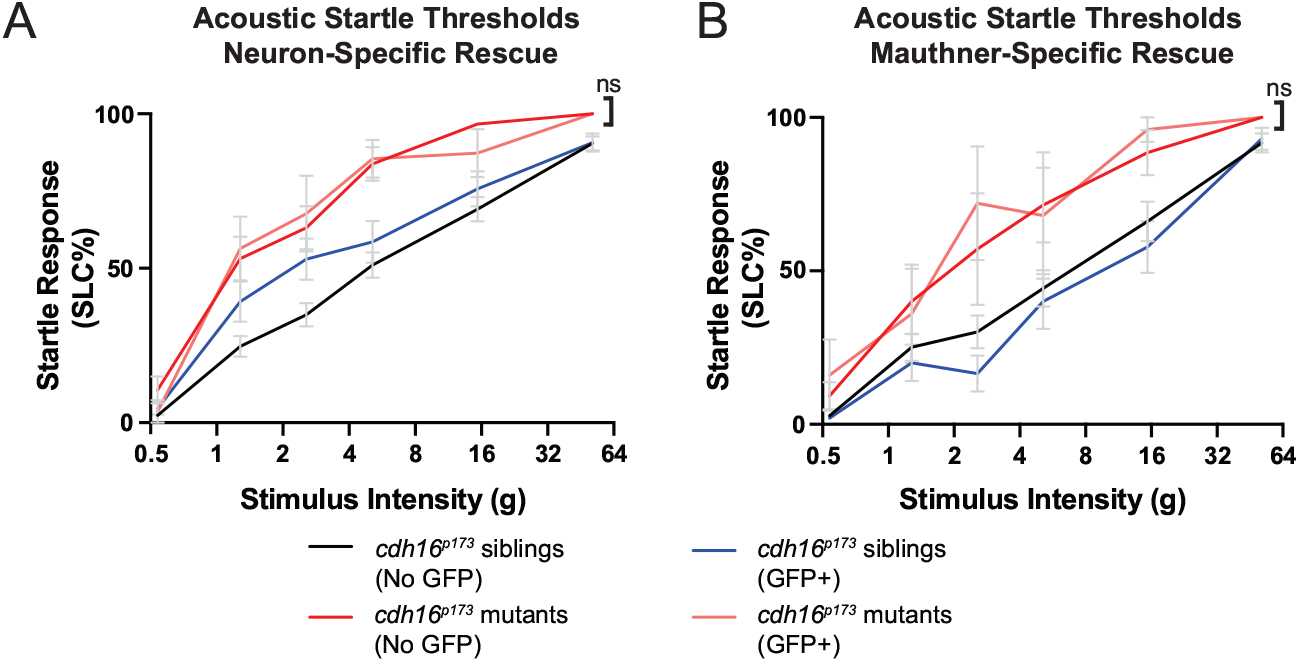
Neuronal expression of *cdh16* does not restore normal startle response thresholds. **A)** Acoustic startle thresholds were measured in 5dpf larvae overexpressing *cdh16* in all neurons. Overexpressing *cdh16-egfp* in mutant neurons (n=11) did not rescue the hypersensitivity phenotype relative to larvae that don’t carry the transgene (n=24) p>0.8 for all stimulus intensities, two-way ANOVA with Tukey’s test for multiple comparisons. Error bars represent SEM. **B**) Acoustic startle thresholds were measured in 5dpf larvae overexpressing *cdh16-egfp* in the Mauthner neuron. Mutants with neuronal overexpression of *cdh16*-egfp (n=5) were no different than those without overexpression (n=7) p>0.8 for all intensities, two-way ANOVA with Tukey’s multiple comparisons test. Error bars represent SEM.

**Supplemental Figure 3.**
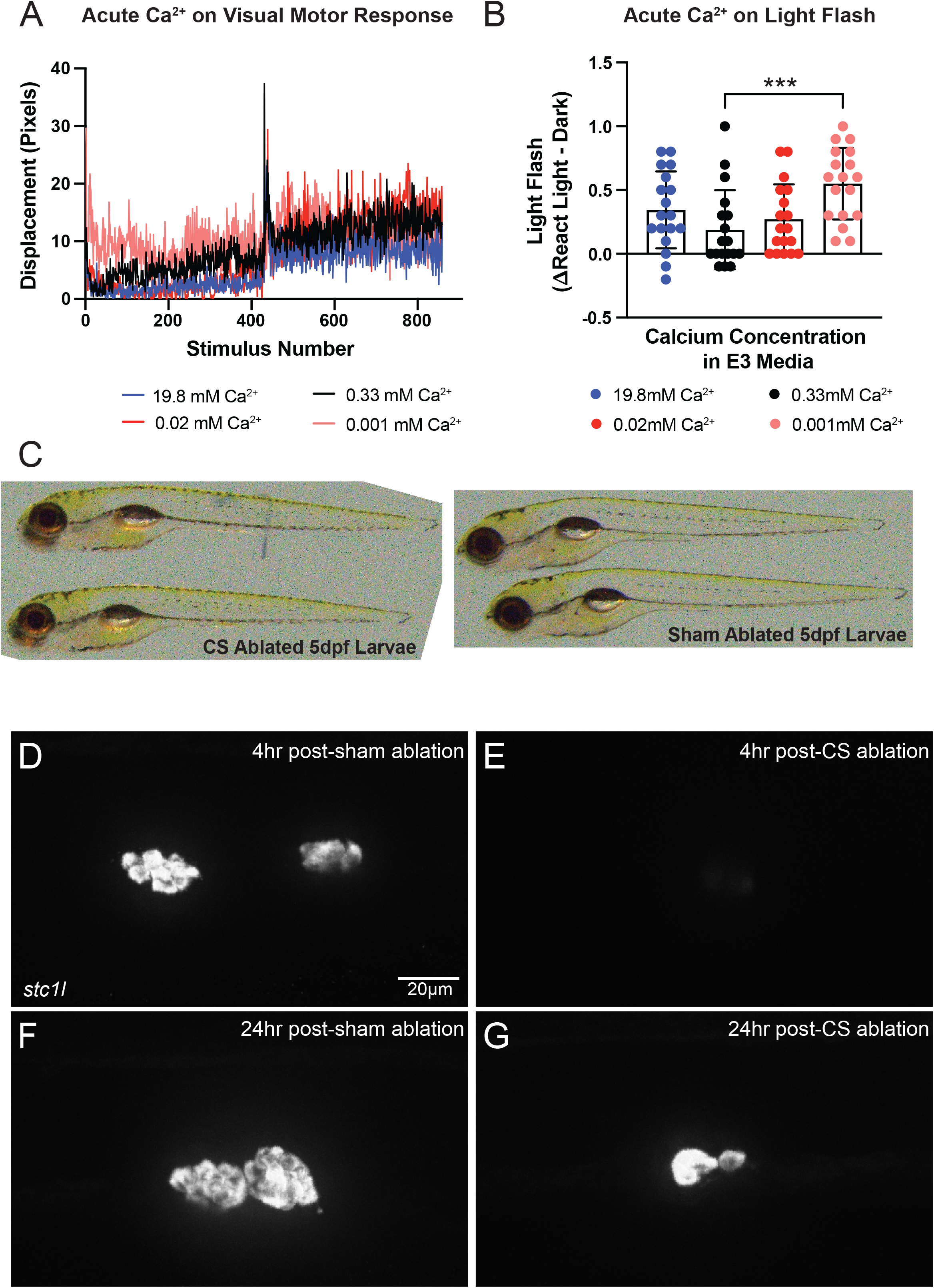
The corpuscles of Stannius (CS) and Ca^2+^ homeostasis are important regulators of behavioral thresholds. **A)** Four calcium (Ca^2+^) concentrations were applied to WT larvae 4 hours before performing behavioral assays at 5dpf. Animals in the lowest (0.001mM) Ca^2+^ concentration (n=18) were more responsive to the lights-on stimulus in the visual motor assay as compared to their siblings in a normal 0.33 mM Ca^2+^ concentration (n=18), p=0.0033, Kruskal-Wallis test with Dunn’s test for multiple comparisons. **B**) Animals in 0.001mM Ca^2+^ (n=18) displayed more robust responses to a light flash than their siblings in 0.33mM Ca^2+^ (n=18) ***p=0.0009, Kruskal-Wallis test with Dunn’s test for multiple comparisons. Error bars represent SD. **C)** Images of 5dpf WT larvae 24 hours after either CS ablation (left) or sham ablation (right). Larvae with ablated corpuscles do not have visible pericardial edema. **(D-G)** *stc1l* HCR to visualize the CS after sham ablation **(D,F)** or CS ablation **(E,G)**. Only a few *stc1l-*positive cells are present in the CS region 4 hours after CS ablation **(E)**, and *stc1l* expression is strongly reduced. By 24 hours post-CS ablation, the structure has partially regenerated **(G)**. Imaged n=10 CS-ablated 4 hours post-ablation, n=5 sham-ablated 4 hours post-ablation, n=10 CS-ablated 24 hours post-ablation, n=5 sham-ablated 24 hours post-ablation.

